# Non-invasive Amelioration of Essential Tremor via Phase-Locked Disruption of its Temporal Coherence

**DOI:** 10.1101/2020.06.23.165498

**Authors:** Sebastian R. Schreglmann, David Wang, Robert Peach, Junheng Li, Xu Zhang, Anna Latorre, Edward Rhodes, Emanuele Panella, Edward S. Boyden, Mauricio Barahona, Sabato Santaniello, Kailash P. Bhatia, John Rothwell, Nir Grossman

**Affiliations:** Institute of Neurology, Department of Clinical and Movement Neuroscience, Queen Square, University College London (UCL), London, WC1N 3BG, UK; Computer Science and Artificial Intelligence Laboratory, Massachussetts Institute of Technology (MIT), Cambridge, MA 02139, USA; NuVu studio Inc, Cambridge, MA 02139, USA; Department of Mathematics and EPSRC Centre for Mathematics of Precision Healthcare, Imperial College London, London, SW7 2AZ, UK; Division of Brain Sciences, Department of Medicine, Imperial College London, London, W12 0NN, UK; UK Dementia Research Institute (UK DRI) at Imperial College London, London, W12 0NN, UK; Biomedical Engineering Department and CT Institute for the Brain and Cognitive Sciences, University of Connecticut, Storrs, CT 06269, USA; Department of Physics, Imperial College London, London, SW7 2AZ, UK; Media Lab, MIT, Cambridge, MA 02139, USA; McGovern Institute for Brain Research, MIT, Cambridge, MA 02139, USA; Broad Institute of Harvard University and MIT, Cambridge, MA 02142, USA; Department of Biological Engineering, MIT, Cambridge, MA 02139, USA; Department of Brain and Cognitive Sciences, MIT, Cambridge, MA 02139, USA; Center for Neurobiological Engineering, MIT, Cambridge, MA 02139, USA; Centre for Bio-Inspired Technology, Department of Electrical and Electronic Engineering, Imperial College London, London, SW7 2AZ London, UK; Centre for Neurotechnology, Imperial College London, London, SW7 2AZ London, UK; Koch Institute for Integrative Cancer Research, MIT, Cambridge, MA 02139, USA; Department of Neurology, Kantonsspital St. Gallen, St.Gallen 9007, Switzerland

## Abstract

Aberrant neural oscillations hallmark numerous brain disorders. Here, we first report a method to track the phase of neural oscillations in real-time via endpoint-corrected Hilbert transform (ecHT) that mitigates the characteristic Gibbs distortion. We then used ecHT to show that the aberrant neural oscillation that hallmarks essential tremor (ET) syndrome, the most common adult movement disorder, can be noninvasively suppressed via electrical stimulation of the cerebellum phase-locked to the tremor. The tremor suppression is sustained after the end of the stimulation and can be phenomenologically predicted. Finally, using feature-based statistical-learning and neurophysiological-modelling we show that the suppression of ET is mechanistically attributed to a disruption of the temporal coherence of the oscillation via perturbation of the tremor generating a cascade of synchronous activity in the olivocerebellar loop. The suppression of aberrant neural oscillation via phase-locked driven disruption of temporal coherence may represent a powerful neuromodulatory strategy to treat brain disorders.

## Introduction

Synchronous oscillatory firing in large populations of neurons has diverse functional roles in the central nervous system (CNS), including regulation of global functional states, endowing connectivity during development, and providing spatiotemporal reference frames for processing of sensory input^1,2^. Aberrant synchronous oscillations have been associated with numerous brain disorders^3,4^. A palpable manifestation of such aberrant oscillation is pathological tremor in essential tremor (ET) syndrome, the most prevalent movement disorder affecting 0.4% of the general population^5^. While the biomolecular origin of ET remains elusive, rendering pharmacological interventions unspecific and often inefficient^6^, its systems-level origin, i.e., oscillatory activity in the cortico-cerebello-thalamo-cortical (CCTC) network, is well established^7^. Invasive systems-level interventions such as lesioning and high-frequency deep brain stimulation (DBS) can successfully treat medication refractory ET^6,8^, but their wide-scale application is limited due to the need for brain surgery. However, such aberrant oscillations fundamentally depend on a delicate cascade of coherent activities across the network components. We here explored whether such a cascade of coherent activities in the CCTC under ET can be disrupted non-invasively by perturbing the synchronous activity of the cerebellum via stimulation that is phase-locked to the tremor oscillation. To phase-lock the stimulation to the tremor oscillation, we first present a strategy to mitigate the Gibbs phenomenon distortion^9^ from the Hilbert transformation^10^ to compute the instantaneous phase of an oscillatory signal in real-time, a strategy that we called endpoint corrected Hilbert transform (ecHT). We then demonstrate that if transcranial alternating current stimulation (tACS) of the cerebellum is phase-locked to ET movement it can suppress its amplitude. Finally, we show that the suppression of ET amplitude is attributed to a disruption of the cascade of coherent activities in the olivocerebellar loop.

## Results

### Real-time computation of instantaneous phase via endpoint corrected Hilbert transform

To enable phase-locking of stimulation to oscillatory activity, we first developed a strategy to compute in real-time the instantaneous phase of oscillatory signals. Traditionally, the instantaneous phase and envelope amplitude, of a band-limited, time-varying oscillatory signal are computed from a complexified version of the signal, known as the analytic signal, in which the real part is the unmodified signal and the imaginary part is the signal’s Hilbert transform^10^. The discrete analytic signal is most accurately and efficiently computed in the frequency domain^11^. However, the Gibbs phenomenon^9^ has made it impossible to accurately compute the instantaneous phase and amplitude at the ends of finite-length analytic signals^12^.

We hypothesized that by applying a causal bandpass filter to the frequency domain representation of the analytic signal we would mitigate the Gibbs phenomenon by establishing a continuity between the two ends of the signal and remove the distortion selectively from the end part of the signal – aka endpoint corrected Hilbert transform (ecHT). See **Online Methods** for a detailed description of the ecHT.

To assess whether the ecHT strategy could effectively mitigate the Gibbs phenomenon at the endpoint of the analytic signal, we computed the Hilbert transform of a test signal, i.e., a finite-length discrete cosine waveform, and quantified the error at the endpoint. **Fig. 1a** and **Fig. 1b** show the Fourier spectra and the Hilbert transforms without the endpoint correction when the signal completed and did not complete full cycles within the sampled time interval, respectively. At the endpoint of the signal without ecHT, the maximal phase error was 179° (mean error 47° ±50° standard deviation, st.d.), and the maximal amplitude error was 191% (76% ±69% st.d.). **Fig. 1c**. **Fig. 1d** shows the same as Fig. 1b but with the endpoint correction. At the endpoint, the ecHT strategy reduced the phase error by at least an order of magnitude (maximal error 12°; mean error 9° ±2° st.d.) and the amplitude error by at least two orders of magnitude (8%; 4% ±2%). The effects of the filter bandwidth and filter order are shown in **Fig. 1f** and **Fig. 1g**, respectively.

**Fig. 1.**
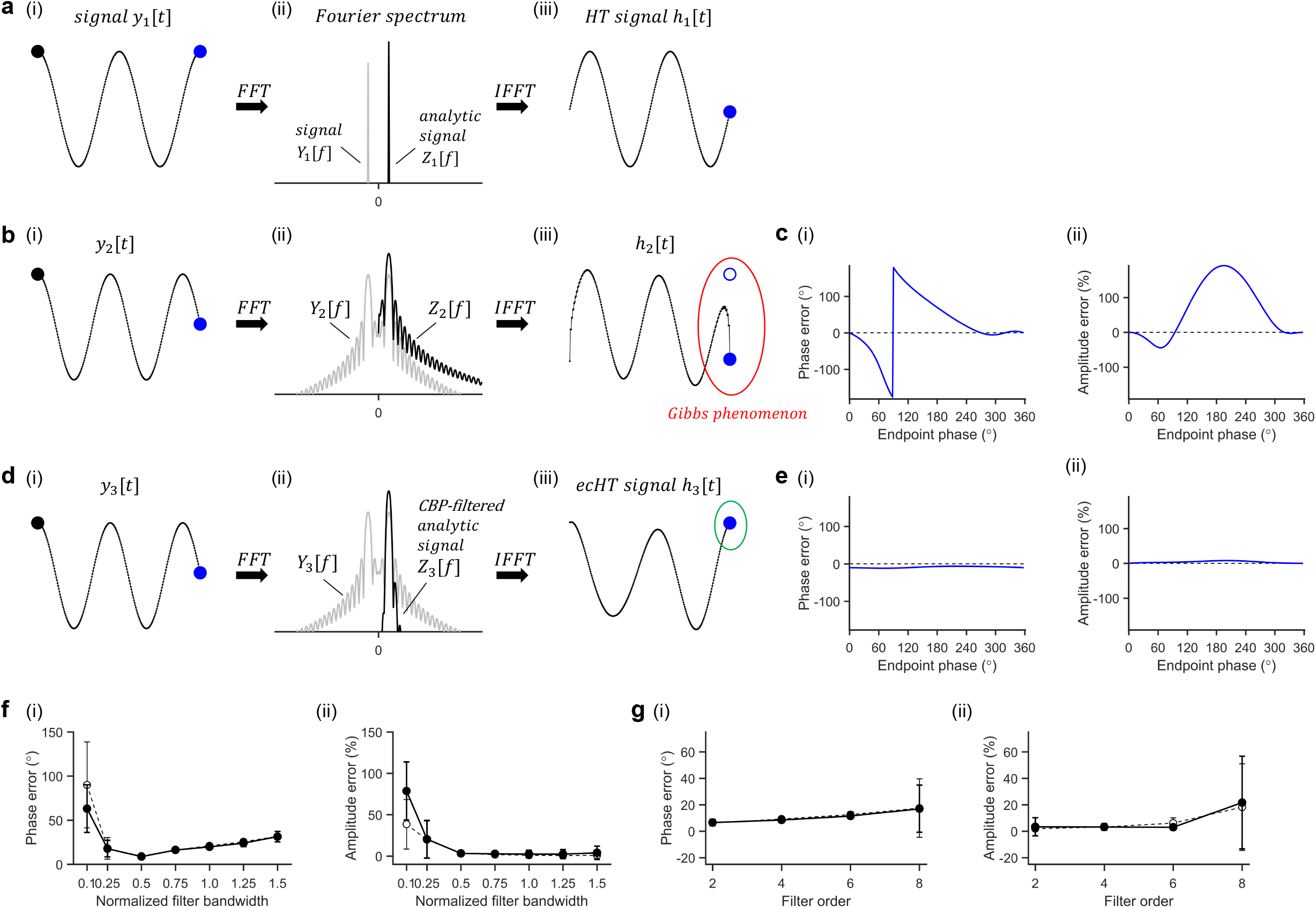
Concept and simulation of real-time computation of instantaneous phase and amplitude via ecHT. **a**, Hilbert transform of a finite, discrete, oscillatory signal completing full cycles. (i) Finite, discrete, oscillatory test signal *y*_1_ - in this example a cosine waveform with an amplitude *A*_1_ = 1, a frequency *f*_1_ = 2 *Hz*, and a phase delay Ø_1_ = 0, sampled at 256 equidistant time points over a finite time interval of 1s, thus completing 2 full cycles over the sampled interval. First data sample and last data sample are marked with black and blue circles, respectively. (ii) Fourier spectrum grey trace, of signal *y*_1_, obtained via fast Fourier transform (FFT) of *y*_1_ (in this example using 256 equidistant frequency points, and the Fourier spectrum *Z*_1_, black trace, of the corresponding analytic signal, obtained from by deleting the negative frequencies and doubling the amplitude of the positive frequencies; y-axis is shown in logscale. *Y*_1_ trace at positive frequencies is overlaid in this plot by the corresponding *Z*_1_ trace. (iii) Hilbert transform *h*_1_ of *y*_1_, i.e., the imaginary part of the analytic signal *z*_1_, obtained via the inverse fast Fourier transform (IFFT) of *Z*_1_ to the same 256 equidistant time points over the finite time interval of 1s. Filled blue circle, computed endpoint; non-filled blue circle, actual endpoint (computed by adding 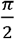 phase lag to *y*_1_; in this case, overlaid by a filled circle). See ^11^ for a detailed description of Hilbert transform computation. **b**, Hilbert transform of a finite, discrete, oscillatory signal not completing full cycles. (i) Finite, discrete, oscillatory test signal *y*_2_ equals to *y*_1_ but with a difference frequency *f*_2_ = 2.25 *Hz*, i.e., completing 2.25 cycles over the 1s sampled interval. First and last data samples are marked as in (ai). (ii) Fourier spectrum *Y*_2_ of the signal *y*_2_, obtained from *y*_2_ as in (aii) and Fourier spectrum *Z*_2_ of the corresponding analytic signal, obtained from *Y*_2_ as in (aii); shown are Fourier spectra sampled using 2048 equidistant frequency points to illustrate the formation of the *sinc* waveform. (iii) Hilbert transform *h*_2_ of *y*_2_, obtained from *Z*_2_ as in (aiii) using the same time sampling as in (aiii). Filled blue circle and non-filled blue circle are computed and actual endpoint as in (aiii); red ellipse, outlines the Gibb phenomenon at the end of the signal. **c**, Computation error of phase and amplitude at the signal’s endpoint obtained via Hilbert transform. (i) Computation error of the instantaneous phase (in °) at the signal’s endpoint for different endpoint phases (in °), simulated by varying *f*_2_ between 2 *Hz* and 3 *Hz* in 0.0056 Hz frequency intervals, quantified as the difference between 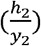 at the last data point and 2*πf*_2_. (ii) Computation error of the instantaneous amplitude (in %) at the signal’s endpoint for different endpoint phases as in (i), quantified as the percentage difference between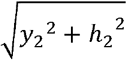 at the last data point and *A*_2_ = 1. **d**, Endpoint corrected Hilbert transformation (ecHT) of a finite, discrete, oscillatory signal not completing full cycles. (**i**) Finite, discrete, oscillatory test signal *y*_3_ equals to *y*_2_. First and last data samples are marked as in (ai). (ii) Fourier spectrum *Y*_3_ of the signal *y*_3_, obtained via FFT of *y_3_* as in (bii), and the Fourier spectrum *Z*_3_ of the corresponding analytic signal, obtained from *Y*_3_ as in (b) but then multiplied by causal bandpass (CBP) filter, in this example, 2nd order Butterworth bandpass filter with a centre frequency *f_3_* and a bandwidth 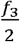. (iii) Hilbert transform *h_3_* of y_3_, obtained from *Z_3_* as in (biii). Filled blue circle and non-filled blue circle are computed and actual endpoint as in (aiii); green ellipse, outlines the mitigation of the Gibb phenomenon at the end of the signal. **e**, Computation error of phase and amplitude at the signal’s endpoint obtained via ecHT. (i) showing the same as (ci) for *y*_3_ and *h_3_*. (ii) showing the same as in (cii) for *y*_3_ and *h*_3_. **f**, Effect of filter’s bandwidth on ecHT computation error at the endpoint. (i) Phase error. (ii) Amplitude error. Shown values are mean ±st.d.; filled black markers, computation error of the instantaneous phase (in °) at the signal’s endpoint obtained as in (ei) but using different filter bandwidths normalized to the filter centre frequency (in this example *f*_3_); nonfilled markers, phase shift (in °) at the same data point, i.e., T, introduced by the filter, obtained by simulating a signal with a longer time interval of 2T (i.e., Gibbs phenomenon is shifted from the T time point). **g**, Effect of filter’s order on ecHT computation error at the endpoint. Showing the same as in (f).

### Cerebellar stimulation phase-locked to essential tremor movement

Next, we deployed the ecHT to test whether stimulation of the cerebellum phase-locked to the tremor movement can perturb ET in a cohort of 11 ET patients (see **Table s1** for demographic details). We measured the tremor movement of the hand, computed its instantaneous phase in real-time, generated eight different stimulating currents – sinusoidal at six different phase lags (0°, 60°, 120°, 180°, 240°, 300°), a control sinusoidal at the tremor frequency without phase-locking, and a sham, and applied them transcranially to the ipsilateral cerebellum via scalp electrodes (mean current amplitude 2.7 ±1 st.d. mA). **Fig. 2a** shows a schematic of the phase-locked stimulation concept, **Fig. 2b** shows a schematic of the electrode configuration and the theoretical distribution of the electric fields in the brain, computed using finite element method (FEM) modelling. **Movie S1** shows a representative video.

**Fi. 2.**
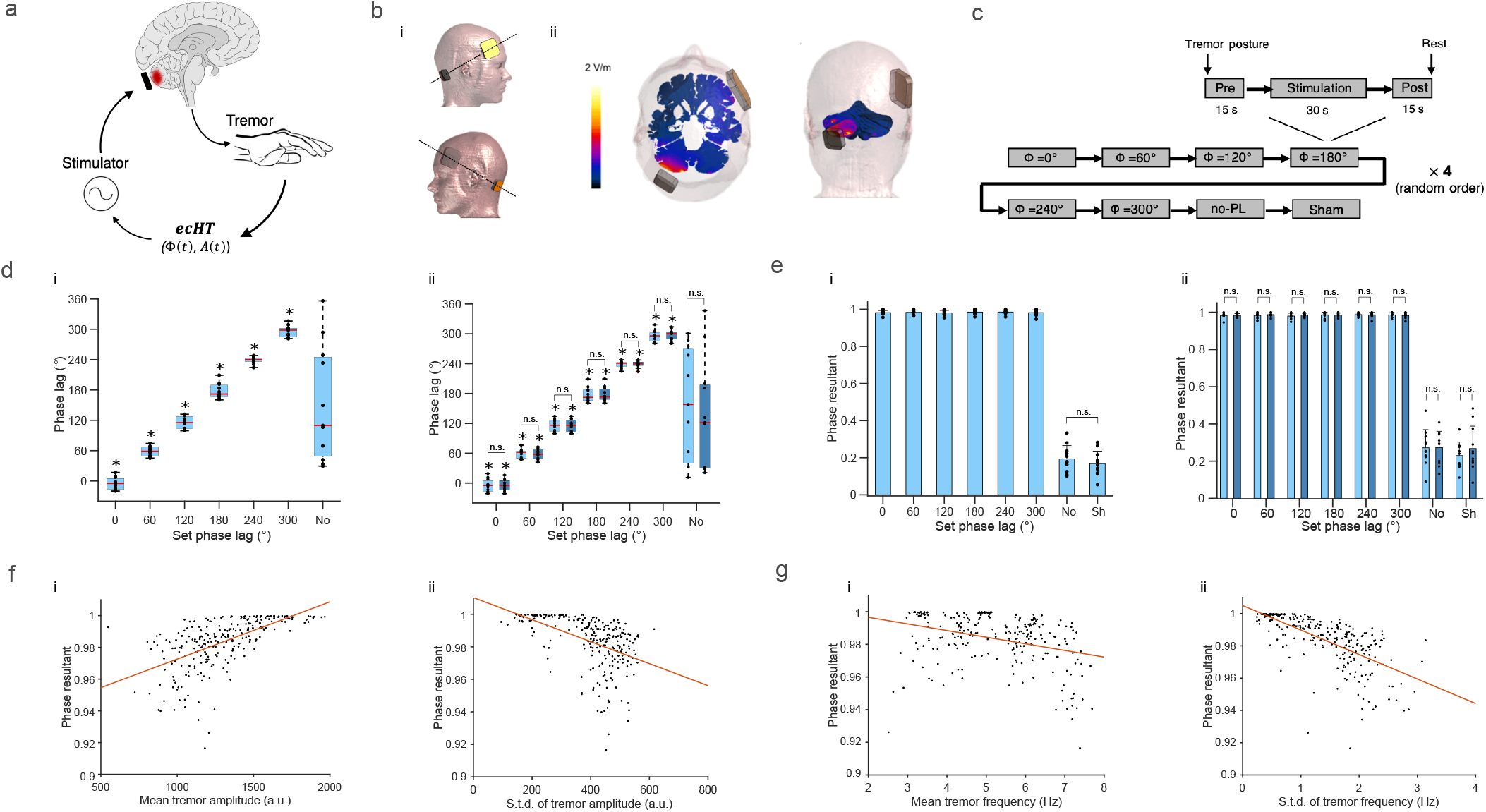
Stimulation of the cerebellum phase-locked to ET movement. **a,** Neuromodulation concept. ET oscillation in the brain is ameliorated by perturbing its pathologic synchrony via cerebellar stimulation that is phase-locked to the hand tremor oscillation. ET oscillation is measured via a motion sensor, instantaneous attributes of the oscillation (i.e., amplitude *A*(*t*), phase *Φ*(*t*)), are computed in real-time using ecHT strategy, stimulating electric currents are delivered to the cerebellum at a fixed phase lag. **b**, Electrode configuration and cerebral electric fields distribution. (i) Stimulating currents were applied via a small (2 x 2 cm^2^) skin electrode placed over the cerebellar hemisphere ipsilateral to measured hand tremor (10% axial nasion-inion distance lateral to inion) and a large (5 x 5 cm^2^) skin electrode placed over the contralateral frontal cortex (between F3-F7 or F4-F8 of the international 10-20 system). (ii) Finite element method (FEM) modelling of induced electric field for a current amplitude of 2 mA. **c**, Schematic of the experimental design. **d**, Phase lag between stimulating currents and tremor movement versus set phase lag during (i) whole stimulation period and (ii) 1st half (light blue) and 2nd half (dark blue) of the stimulation period. ‘No’, control sinusoidal current at the tremor frequency but without phaselocking; shown are box, 25% and 75% percentile values; horizontal red line, median value; horizontal black lines, data range; black markers, patients’ values; ‘No’, stimulation with no phase locking; * p < 0.05, Omnibus test. See **Table S2** for between conditions statistics. **e**, Mean phase resultant vector length versus set phase lag during the same periods as in (d); shown are mean ±st.d.; markers show patients’ values; ‘No’, stimulation with no phase locking; ‘Sh’, sham stimulation. See **Table S3** for full statistics. **f**, Mean phase resultant vector length versus (i) tremor amplitude and (ii) st.d. of tremor amplitude; shown black markers are trials’ mean values. Red line is the linear regression fit, in (i) line slope m=0.59, p<10^−5^, Pearson correlation test, (ii) m=-0.49, p<10^−16^. **g**, Mean phase resultant vector length versus (i) tremor frequency and (ii) st.d. of tremor frequency, shown as in (f); linear regression fit in (i) m=-0.33, p<10^−7^, (ii) m=-0.66, p<10^−32^.

We applied each stimulation condition in a block of 60s during which the patients maintained a tremor evoking posture. Each block consisted of a 30s stimulation period (including 5s of ramp-up and 5s of ramp-down) and 15s stimulation-free periods before and after. We repeated the stimulation conditions four times in a double-blinded random order with a 30s rest interval between conditions and 5-10min rest interval between sessions of eight stimulation conditions (see **Fig. 2c** for a schematic of the study design and **Online Methods**).

To assess whether the stimulating currents were delivered at accurate phase-lag, we computed, offline using Hilbert transform, the lag between the instantaneous phase of the stimulation waveforms and the instantaneous phase of the tremor movement waveforms. We found that during the phase-locked stimulation, the phase-lag distribution of each condition was narrow and different from the other conditions throughout the stimulation period (**Fig. 2d(i)**) and during the first and second halve periods (**Fig. 2d(ii)**), (p<10^−8^ for all periods; Fisher test; see **Table S2** for full statistics). The difference between the measured phase-lag and the set phase-lag was small, i.e., 3° ±11° (mean ±st.d), across all the phase-locked conditions. The mean resultant vector length (quantifying the circular spread)^13^, was close to one, i.e., 0.98 ±0.01, across all the conditions, and did not differ between conditions throughout the stimulation period (**Fig. 2e(i)**), and during the first and second halve periods (**Fig. 2e(ii)**; p>0.95**_**for all periods; one-way ANOVA, see **Table S3** for full statistics). The mean resultant vector length was slightly larger at stimulation blocks with higher tremor amplitude (**Fig. 2f(i)**) and was slightly smaller at stimulation blocks with higher tremor amplitude st.d. (**Fig. 2f(ii)**) higher tremor frequency (**Fig. 2g(i)**) and higher tremor frequency st.d. (**Fig. 2g(ii)**). In contrast, during the sinusoidal stimulation without phase-locking, the phase-lag distribution was not different from a uniform distribution (**Fig. 2d(i-ii);** p>0.4 for all periods; Omnibus test). The mean resultant vector length was small, i.e., 0.19 ±0.071, and did not differ from sham stimulation (p=0.37, paired Wilcoxon signed-rank test), indicating that the stimulation did not entrain the tremor phase (**Fig. 2d-e** and **Table S3**). Across all stimulation conditions, the mean resultant vector length was not different in trials in which patients reported sensation underneath the electrodes and trials in which no sensation was reported (p=0.3, Paired sign-rank test).

### Phase-dependent suppression of essential tremor amplitude

After establishing that the stimulating currents were delivered at the desired phase lags, we assessed whether they affected the tremor amplitude. To quantify the stimulation effect relative to the baseline period and relative to the effect of sham stimulation, we computed, for each patient, the z-score of the tremor amplitude relative to the mean and the st.d. of the tremor amplitude during baseline in each stimulation condition, and then subtracted the median z-score of the tremor amplitude during sham stimulation (the tremor frequency and amplitude were not significantly different, see **Table S1** for full statistical details). To examine the temporal dynamics of the effect we quantified the z-score values during the first half and second half of the stimulation period, as well as during the post-stimulation period.

We found that the stimulation at the tremor frequency without phase-locking resulted in a tremor amplitude reduction, yet not statistically significant (**Fig. 3a**). A significant tremor amplitude change (reduction or increase) occurred in only a small number of patients (**Fig. 3b**). Across these subsets of patients, the change was statistically significant only in those showing a reduction and only during the first half of the stimulation (**Fig. 3c-d**). In contrast, stimulation that was phase-locked to the tremor movement resulted in a significant reduction in the tremor amplitude, that increased throughout the stimulation period and sustained during the post stimulation period, **Fig. 3e**. The number of patients who showed a significant reduction in the tremor amplitude was significant during the second half of the stimulation and the post-stimulation period, while the number of patients who showed a significant increase in the tremor amplitude was not significant throughout, **Fig. 3f** (p-value threshold of amplitude change was Bonferroni corrected for six phase-locked conditions). Across these subsets of patients, the reduction/increase in the tremor amplitude was statistically significant throughout (**Fig. 3g-h**). The change in tremor amplitude was not different between sessions (p=0.64, ANOVA; p=0.32, linear mixed effect model with sessions as a predictor variable). Across all stimulation conditions, the z-score tremor amplitude was not different in trials in which patients reported sensation underneath the electrodes and trials in which no sensation was reported (p=0.54, paired t-test).

**Fig. 3.**
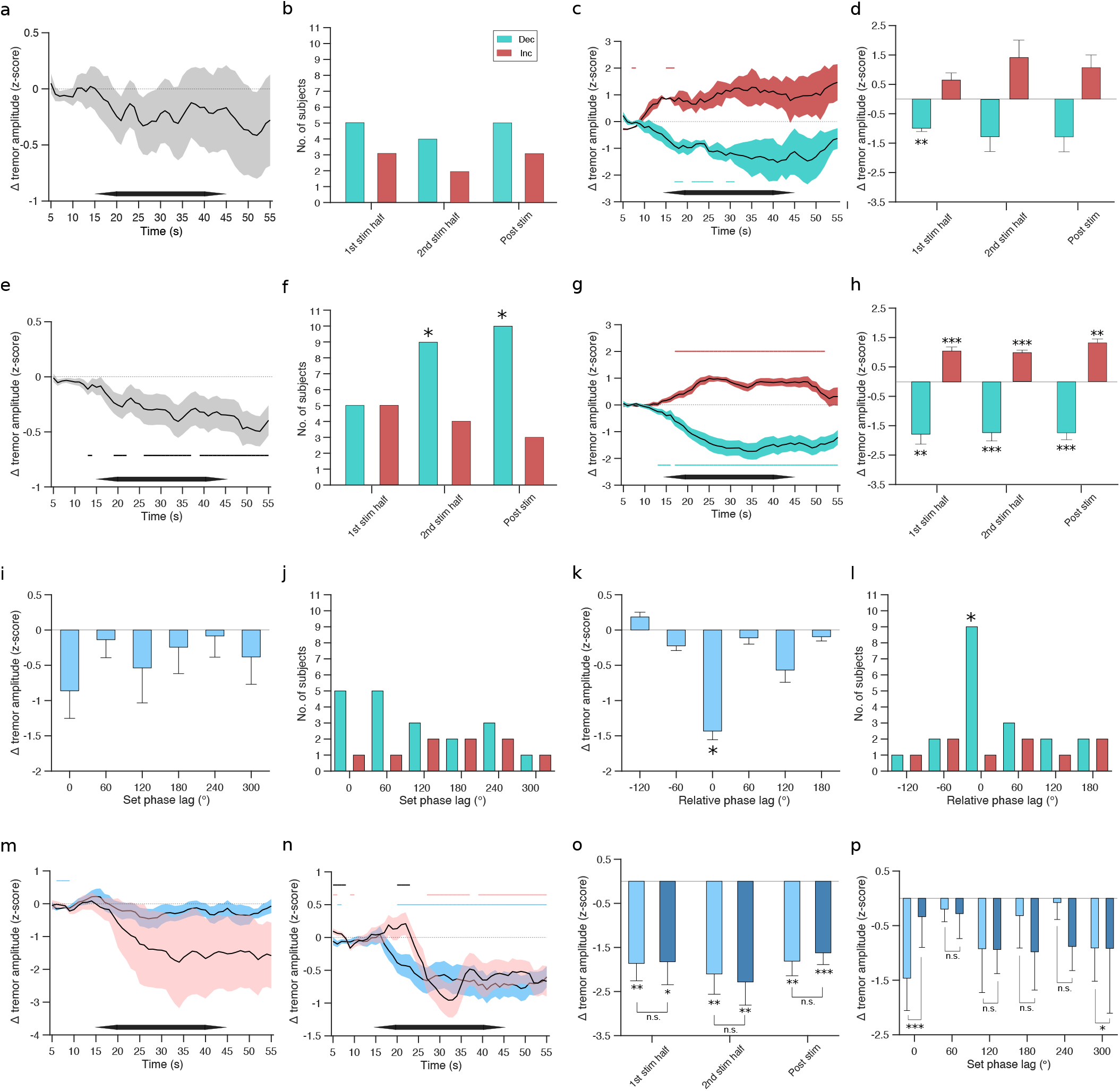
Characterization of change in tremor amplitude induced by stimulation. Tremor amplitudes were z-scored relative to baseline, by subtracting the mean baseline amplitude and dividing by the st.d. of baseline amplitude, and were expressed relative to sham stimulation, by subtracting the median z-score during sham. **a**-**d**, Stimulating currents were applied at the tremor frequency but without phase-locking. **a**, Change in tremor amplitude over time, shown are mean ±s.e.m. (standard error of the mean) z-score computed using 10 s window every 1 s between 5 s and 55 s from block onset (baseline period used, 3 s to 13 s from block onset); horizontal black bar outlines stimulation period. **b**, Number of patients with a statistically significant reduction (turquoise bars) and increase (red bars) in tremor amplitude during the first half of stimulation period (‘1st stim half’), the second half of stimulation period (‘2nd stim half’), and during the post-stimulation period (‘post stim’); reduction: 1st stim half, 5 patients showed amplitude reduction, 3 patients showed no amplitude change, p=0.66; 2nd stim half, 4, 5, p=1; post stim, 5, 3, p=0.66; Fisher exact test. Increase: 3, 5, p=1; 2, 5, p=0.36; 3, 3, p=1. **c**, Change in tremor amplitude over time across patients with a significant reduction (turquoise) and increase (red) in tremor amplitude during 2nd stim half in (b), shown are mean ±s.e.m. z-score; horizontal turquoise and red lines show corresponding epochs with significant z-score amplitude; horizontal black bar outlines stimulation period. **d**, Change in tremor amplitude during 1st stim half, 2nd stim half, and the post stim periods in (c), shown are mean ± s,e.m.. **e**-**l**, Stimulating currents were phase-locked to the tremor movement. **e**, Change in tremor amplitude over time, showing the same as in (a); horizontal black lines show epochs with significant z-score amplitude. **f**, Number of patients with statistically significant reduction and increase in tremor amplitude in (e), showing the same as in (b); decrease: 5, 1, p=0.15; 9, 1, p<0.005; 10, 0, p<0.00005; increase: 5, 1, p=0.15; 4, 1, p=0.31; 3, 0, p=0.21. Here, 3 patients during the 2nd stim half period and 2 patients during the post-stim period showed a significant reduction in one phase-lag and a significant increase in another phase-lag. In addition, 3 patients during the 2nd stim half period and 2 patients during the post-stim period showed also a significant reduction during stimulation without phase-locking, however the reduction was mostly smaller without phase-locking (see Table S4 for full statistical details). **g**, Change in tremor amplitude over time across patients with decreased and increased tremor amplitude during 2nd stim half in (f), showing the same as (c). **h**, Change in tremor amplitude in (g), showing the same as in (d). **i**-**l**, Effect of the phase lag of stimulation. Shown values are for 2nd stim half. See **Table S4** for complete statistical data including 1st stim half and post-stimulation period. **i**, Change in tremor amplitude versus stimulation phase lag, shown are mean ± s.e.m.. **j**, Number of patients with reduced (turquoise bars) and increased (red bars) tremor amplitude during 2nd stim half versus stimulation phase lag. **k**, Same as (i), i.e., with all patients and phase lags, but phase lags of each patient are expressed relative to the phase lag showing the largest reduction in tremor amplitude and wrap to ±180°. **l**, Same as (j) but phase lags of each patient are expressed as in (k). **m-p**, Characterization of tremor amplitude during the repeated experiment in a subset of patients (patients 1,2,3,6, 9 and 11), see **Table S5** for phase-lag statistics. **m**, Change in tremor amplitude over time when stimulating currents were applied at the tremor frequency but without phase-locking, showing original experiment (blue) and repeated experiment (red); horizontal blue and red lines show epochs with significant z-score amplitude in original and repeated experiments, respectively; horizontal black lines show epochs with a significant difference in z-score amplitude between original and repeated experiments. **n**, Same as (m) but stimulating currents were phase-locked to the tremor movement. **o**, Change in tremor amplitude during 1st stim half, 2nd stim half, and the post-stim periods in (n), shown are mean ± s,e.m. in original experiment (light blue) and repeated experiment (dark blue); see **Table S6** for full statistics. **p**, Change in tremor amplitude versus stimulation phase lag, colour scheme as in (o); see **Table S7** for full statistics. Significance of z-score amplitude in (a), (c), (d), (e), (g), (h), (i), (k), and (m)-(p) was analysed using unpaired t-test, p-value threshold in (g) was Bonferroni corrected for multiple comparisons of stimulation conditions; Significance in (k) was also analysed using 2-sample Kolmogorov-Smirnov test (again Bonferroni corrected for multiple comparisons of stimulation conditions) against a surrogate distribution of z-score values (−120°, p=0.05; −60°, p=0.1; 0°, p=0.002; 60°, p=0.09; 120°, p=0.09; 180°, p=0.1). Significance of z-score amplitude difference between original and repeated experiments in (m-p) was analysed using paired t-test. * indicates p < 0.05, ** indicates p<0.005, *** indicates p<0.00005, n.s. indicates non-significant across the figure. Significance of the number of patients who showed a significant increase/decrease in the z-score tremor amplitude in (b), (f), (j) and (l) was analysed using Fisher exact test against the number of patients who did not show a change in the z-score tremor amplitude; * indicates p < 0.05 for a larger number of patients with increase/decrease in z-score tremor amplitude.

Comparing the phase-locked conditions, we found that the reduction in the tremor amplitude was close to significance (not corrected) only at a phase-lag of 0° (**Fig. 3i**) but the number of patients who showed a significant reduction in the tremor amplitude was not significant (**Fig. 3j**). However, if the phase lags of individual patients were expressed relative to the phase lag that resulted in the largest reduction of their tremor amplitude, the reduction in the tremor amplitude and the number of patients who showed a significant reduction, were statistically significant □ indicating a narrow range of efficacious phase that can vary between patients (**Fig. 3k-l**, see **Table S4** for complete statistical details).

To test whether the effect of the stimulation on the tremor amplitude is reproducible, we repeated the experiment in a subset of patients (n=6, including patients 1,2,3,6, and 11 who showed a reduction in the tremor amplitude and patient 9 who did not; see **Table s1** for demographic and clinical details during the repeated experiment) and analysed the data in the same way as in the original experiment. We found that in the repetition experiment the stimulation currents were delivered at the same phase-lag accuracy as in the original experiment (**Table S5**). As before, stimulation at the tremor frequency without phase-locking resulted in a tremor amplitude reduction, yet not statistically significant (**Fig. 3m**), however stimulation currents that were phase-locked to the tremor movement resulted in a significant reduction in the tremor amplitude that was sustained during the post-stimulation period (**Fig. 3n**). The patients who showed a significant reduction in the tremor amplitude during the stimulation period in the original experiment also showed a significant reduction in the tremor amplitude in the repetition experiment (see **Table S6** for full statistics). The z-score reduction in the tremor amplitude across those patients was not different from the original experiment (**Fig. 3o**). Comparing the phase-locked conditions, we found that at across the cohort the reduction in the tremor amplitude was smaller at phase-lag of 0° and larger at phase-lag of 300° (**Fig. 3p**, see also **Table S7** for full statistics). Within individual patients the phase-lag values that reduced the tremor amplitude were consistent in only 20% of the cases.

### Prediction of patients’ response from distinct features of the tremor movement

Next, we sought to explore whether the variability in the patients’ response to the stimulation can be attributed to certain characteristics of their ET condition. We divided the patients into two groups, i.e., a ‘responder’ group (n=7, including patients 1,2,3,6,8,9, and 11) and a ‘nonresponder’ group (n=4, patients 4,5,7, and 10) - a patient was defined a ‘responder’ if his/her tremor amplitude decreased in at least one of the tested stimulation phases relative to sham and did not increase in any of the tested stimulation phases relative to sham, and a ‘non-responders’ if his/her tremor amplitude increased in at least one of the tested stimulation phases relative to sham or did not change in any of the tested stimulation phases relative to sham.

We first assessed whether certain clinical or demographic characteristics can distinguish between responder and non-responder groups but found only non-significant trends of younger age (p=0.07, Wilcoxon rank-sum test) and higher tremor frequency (p=0.08) in responders (see **Table S1** for full statistical details). In addition, we did not find a difference between the groups in the amplitude of the applied currents (p = 0.8).

We then explored whether certain characteristics of the tremor movement can distinguish between the two groups. We deployed a feature-based statistical learning strategy^14^ to extract 7873 different time-series features from a 10s segment of the tremor movement before the onset of the stimulation in all the trials with phase-locked stimulation (301 trials in total, including 28 trials per patient except patient 3 in which only 21 trials were recorded); exemplary tremor traces are shown in **Fig. 4a**. We then used the features and a support vector machine (SVM) with a linear kernel to classify the tremor trials according to the subjects’ responsiveness to a phase-locked stimulation. We found that using all the features, the tremor trials could be classified according to the patients’ response with an accuracy of 97% (F-score of 96). However, even a small number of features was sufficient for high accuracy classification, using the top 1, 5, 10, and 40 features with highest single-feature classification accuracy, the tremor trials could be classified with an accuracy of 83%, 81%, 86%, and 92% (F-score of 82, 80, 85, and 91), respectively (**Fig. 4b**).

**Fig. 4.**
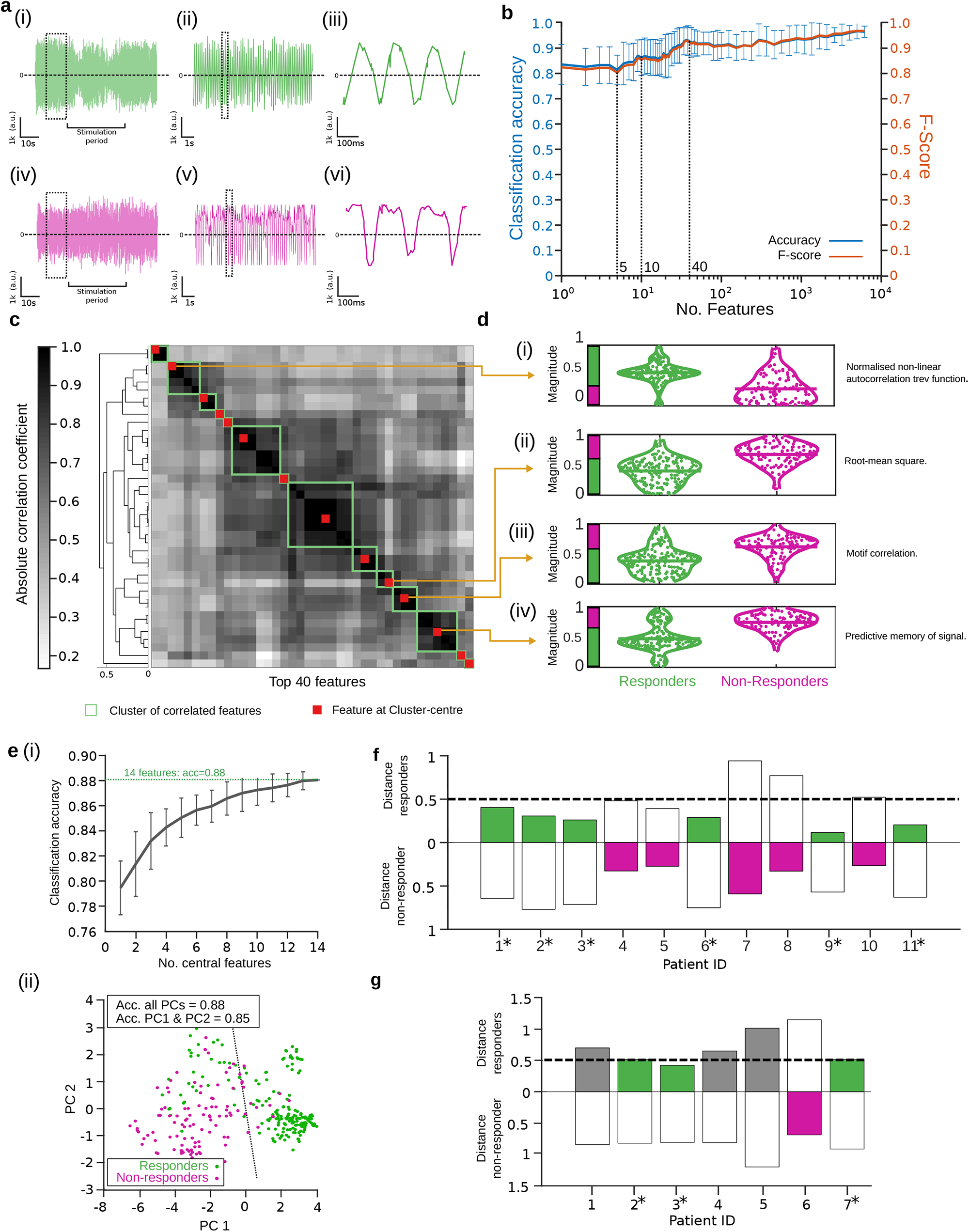
Classification and prediction of patient’s response via features extraction and statistical learning of the tremor movement. Patients were divided into ‘responders’ and ‘non-responders’ groups. Normalized magnitude of 7873 different features were extracted from a 10s baseline period (i.e., before the onset of the stimulation) of the tremor time-series traces of blocks with phase-locked stimulation and were then used to classify the tremor traces according to the patient’s response to the stimulation via supervised statistical learning. **a**, Exemplary recording of tremor movement from a patient that showed a reduction in tremor amplitude during phase-locked stimulation (i-iii) and from a patient that did not show a reduction in tremor amplitude during the stimulation (iv-vi), relative to sham. **b**, Classification accuracy (blue) and F-score (orange; quantifying both precision and recall) of patients’ response as a function of the number of features. Shown are mean and st.d. values of the 10-fold cross-validation. **c**, Most informative features of the class structure. Shown are the 40 top predictive features in **b**, clustered according to correlation coefficient and re-ordered according to the clustering; green box, outline of a feature cluster; red square, central feature of a cluster. See **Table S8** for a list of the features at the cluster’s centre. Feature values of individual patients did not differ between trials (p>0.5; ANOVA). **d**, Normalized magnitude of exemplary features shown in **c** at the centres of the clusters of correlated features. Green, ‘responders’ patients; magenta, ‘non-responders’ patients. ‘Normalised non-linear autocorrelation trev function’ (feature ID, 1129 in ^14^), estimates the amplitude symmetry of the periodic signal relative to zero using autocorrelation; ‘Root mean square’ (feature ID, 16), estimates the amplitude of the periodic signal using root mean square; ‘Motif correlation’ (feature ID, 3561), estimates the time symmetry of the periodic signal (i.e., relative period of signal with an amplitude above and below average); ‘Predictive memory of signal’ (feature ID 2390), estimates the variance in the predictability of a data-point within the periodic signal from the previous data-points. See Fulcher et al.^14^ for a more detailed description of the features. **e**, Classification accuracy of patients’ response using the 14 most informative features, i.e., the features shown in **c** at the centres of the clusters of correlated features, showing (i) mean classification accuracy ±st.d. vs number of features, each repeated 100 times with a random selection of features out of the 14 most informative features, and (ii) 2D principal component analysis (PCA) plots of classification using all 14 features. Acc, classification accuracy; PC, principal component. **f**, Euclidean distance between feature centroids of individual patients and the feature centroids of the responders’ and non-responders’ classes, using the 14 most informative features (distance of responders to responders’ class, mean 0.35 ±0.2 st.d.; responders to non-responders class, 0.6 ±0.25; non-responders and responders class, 0.65 ±0.15; non-responders and non-responders class, 0.35 ±0.15); *, responding patient; green bar, distance to responders class < 0.5 & distance to responders class < distance to nonresponders class; magenta bar, distance to responders class > distance to non-responders class. **g**, Same as f but for a new cohort of patients, showing distances to the same centroids of responders’ and non-responders’ classes in f, i.e., of the original patient cohort; grey bar, distance to responders class < 0.5 but distance to responders class > distance to non-responders class. See **Table S10** for phase-locking and **Table S11** tremor amplitude statistics of this cohort.

We then used a hierarchical cluster tree approach to search for the most informative features among the 40 features with the highest classification accuracy (**Fig. 4c**). We identified 14 clusters of correlated features and extracted the corresponding features at the centre of those clusters – the list of the most informative features is given in **Table S8** and the magnitude probability density plots of exemplary features are shown in **Fig. 4d** (the classification accuracy plateaued at approximately 14 features, **Fig. 4e**). The extracted features revealed that the tremor movement in responders was smaller (**Fig. 4dii**), had a more sinusoidal like regularity (**Fig. diii** and **Fig. div**), and had a higher amplitude symmetry relative to zero (**Fig. 4di**). The Euclidean distance between feature centroids of the responders class and non-responders class was 0.55 (feature centroid of a class was computed by averaging the features across the corresponding samples). The feature centroids of individual patients who responded to the stimulation located at a distance <0.5 to the feature centroid of the responders class and had a longer distance to the feature centroid of the non-responders class (exception was patient 8; **Fig. 4f**).

To test whether these features of the tremor movement can potentially help to predict the response of patients to the stimulation, we repeated the experiment in a new cohort of seven ET patients. We analysed the data in the same way as in the original cohort and extracted the same 14 features from the 10s tremor movement before the stimulation onset (see **Table S9** for demographic details, see **Table S10** for phase-locking and **Table S11** tremor amplitude statistics). We found that three patients (i.e., patients 2,3, and 7) responded to the stimulation based on the aforementioned responding criterion. The feature centroids of these patients, but not the rest of the cohort, were located at ≤0.5 distance to the feature centroid of the responders class from the original cohort and had a longer distance to the feature centroid of the non-responders class from that cohort (**Fig. 4g**) indicating a consistency in the relationship between the features of the tremor movement and the response to the stimulation.

### Suppression of essential tremor amplitude is underpinned by disruption of temporal coherence of movement

After establishing that patients who responded to stimulation had distinct characteristics of tremor movement during baseline, we next sought to explore whether the change in tremor amplitude during stimulation was associated with a change in other characteristics of tremor movement. We divided all the tremor trials with phase-locked stimulation (again 301 trials in total) into three datasets according to the change in tremor amplitude during stimulation relative to sham, i.e., trials with a decrease in tremor amplitude (‘decrease’; 58 trials from 11 subjects), trials with an increase in tremor amplitude (‘increase’; 51 trials from 10 subjects; patient 6, did not show an increase in tremor amplitude in any phase-locked condition), and trials without a change in tremor amplitude (‘no-change’; 192 trials from 11 subjects).

We then deployed the same feature-based statistical learning strategy^14^ to test whether the characteristics of the tremor movement can distinguish between the stimulation and baseline periods in these three datasets. We extracted the same 7873 features as before from a 10s segment of the tremor movement before the onset of the stimulation and from a corresponding 10s segment during the middle of the stimulation; exemplary tremor traces with tremor amplitude ‘decrease’ and ‘increase’ are shown in **Fig. 5a** and **Fig. 5b**, respectively. We then used the features and the same SVM as before to classify the tremor trials according to the period class, i.e., ‘baseline’, or ‘stimulation’. We found that the ‘decrease’ dataset had a higher probability of classification with high accuracy compared to the ‘increase’ and the ‘no-change’ datasets (**Fig. 5c**).

**Fig. 5.**
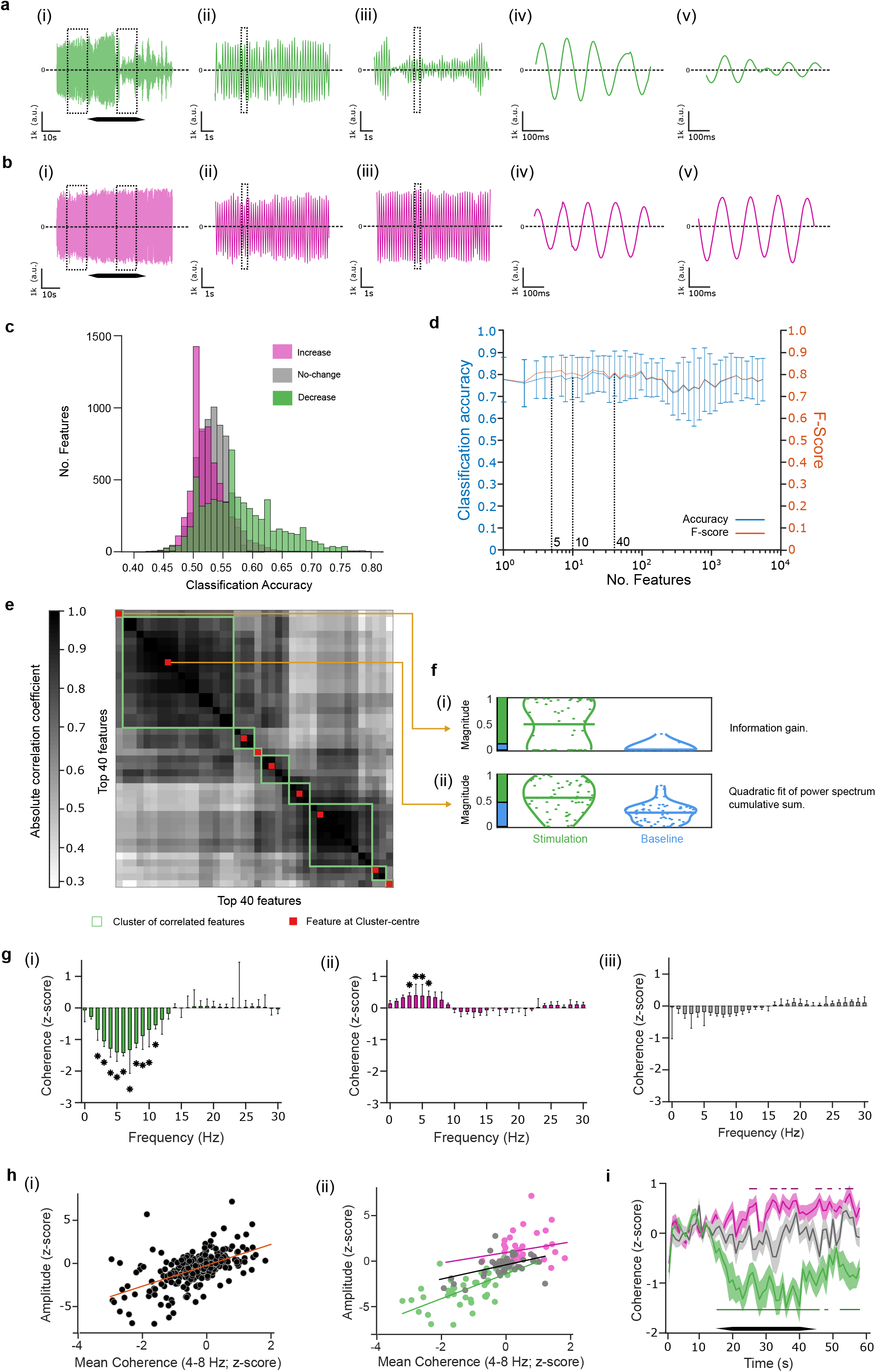
Change in ET amplitude is linked to change in temporal coherence of the tremor movement. Recorded traces were divided into ‘decrease’, ‘increase’, and ‘no-change’ datasets according to the effect of the stimulation on the tremor amplitude. Normalized magnitude of the same 7873 features as in **Fig. 4** were extracted from 10s during stimulation period (‘stimulation’ class) and from 10s during the corresponding pre-stimulation period (‘baseline’ class) and were then used to classify the tremor traces via supervised statistical learning according to the period class. **a**, Exemplary recording of tremor movement from a patient (patient 1) during stimulation at a phase that resulted in a reduction of tremor amplitude relative to sham. (i) full 60s recording including 15s baseline period, 30s stimulation period (indicated with black hexagon; including 5s ramp-up and 5s ramp-down), and 15s poststimulation period. (ii) and (iii) show magnified view the 10s baseline trace period and the 10s stimulation period indicated by a box in (i); (iv) and (v) show magnified view of the trace region indicated by a box in (ii) and (iii), respectively. **b**, Exemplary recording of tremor movement from the same patient as in (a) but during stimulation at a phase that resulted in a small increase of tremor amplitude. (i-v) as in (a). **c**, Probability distribution histogram of the feature-based classification accuracy according to the period class (i.e., ‘baseline’ and ‘stimulation’) of the ‘decrease’ (green), the ‘increase’ (magenta), and the ‘no-change’ (grey) datasets. Significance was tested using pairwise Kolmogorov-Smirnov test ‘decrease’ vs. ‘increase’, p=0.01; ‘decrease’ vs. ‘no-change’, p=0.008; ‘increase’ vs. ‘no-change’, p =0.45; and against a null distribution, generated by assigning random values to the feature), ‘decrease’, p=0.005; ‘increase’, p=0.34; ‘no-change’, p=0.58. **d**, Classification accuracy (blue) and F-score (orange) of the time-series traces in the ‘decrease’ dataset according to the period class (i.e., ‘baseline’ and ‘stimulation’) as a function of the number of features. Shown are mean and st.d. values of the 10-fold cross-validation. **e**, Most informative features for the class structure in the ‘decrease’ dataset. Shown are the 40 top predictive features in (d), clustered according to correlation coefficient and re-ordered according to the clustering; green box, outline of a feature cluster; red square, central feature of a cluster. See **Table S7** for a list of the features at the cluster’s centre. **f**, Normalized magnitude of features shown in (e) at the centres of the clusters of correlated features. Green, ‘stimulation’ period; blue, ‘baseline’ period. ‘Information gain’ (feature ID, 2315 in ^14^), estimates the gain in a data-point relative to the previous data-points; ‘Quadratic fit of power spectrum cumulative sum’ (feature ID, 4432), computes the accuracy, i.e., r square value, of fitting the cumulative sum of the power spectrum with a quadratic function. See Fulcher et al. ^14^ for a more detailed description of the features. **g**, Change in tremor’s temporal coherence, quantified by computing the magnitude-squared coherence across 1s epochs during stimulation and during baseline. Shown values are mean ± st.d. z-score during stimulation period relative to mean and st.d. of baseline period from (i) ‘decrease’ dataset, (ii) ‘increase’ dataset, and (iii) dataset of sham stimulation (‘sham’); * p<0.0005; significance was assessed using unpaired t-test with Bonferroni corrections for multiple comparisons of frequency bins and datasets. **h**, Correlation between change in tremor’s amplitude and change in tremor’s temporal coherence at the tremor frequency band (i.e., 4-8 Hz). Each datapoint is z-score of tremor’s amplitude versus z-score of tremor’s temporal coherence from a single trial during stimulation period relative to the baseline period (computed as in (g)) from (i) the combined datasets of ‘decrease’, ‘increase’ and ‘sham’ and (ii) the individual datasets with ‘decrease’, green; ‘increase’, magenta; ‘sham’, grey. Pearson correlation test values: combined dataset, c=0.2, m=1.2, R^2^ =0.32; ‘decrease’ dataset, c=-1.4, m=1.35, R^2^ =0.49; ‘increase’ dataset, c=0.94, m=0.58, R^2^ =0.004; ‘sham’ dataset, c=-0.3, m=0.78, R^2^ =0.32 (c, line y-intercept; m, line slope). **i**, Change in tremor’s temporal coherence at the tremor frequency band (same as in (h)) over time, quantified by computing the magnitude-squared coherence between each 1s epoch and its preceding epoch during stimulation and during baseline. Shown values are mean ± st.d. z-score at each epoch during the stimulation period relative to the mean and st.d. across epochs during the baseline period from the same ‘decrease’ (green), ‘increase’ (magenta) and ‘sham’ (grey) datasets as in (g); horizontal lines show epochs with significant z-score temporal coherence, characterized using unpaired t-test with Bonferroni corrections for multiple comparisons of datasets; horizontal black bar outlines stimulation period.

Focusing on the ‘decrease’ dataset, we found that using all the features, the tremor trials during stimulation and baseline could be classified with an accuracy of 79% (F-score of 79). However, the classification accuracy was dominated by only a few features, using the top 1, 5, 10, and 40 features with highest single-feature classification accuracy, the tremor trials could be classified with an accuracy of 78%, 79%, 79%, and 80% (F-score of 78, 81, 81, and 81, respectively; **Fig. 5d**). We then used, as before, the hierarchical cluster tree approach with a between-feature correlation threshold of 0.2 to search for the most informative features among the 40 features with the highest classification accuracy (**Fig. 5e**). We identified 9 clusters of correlated features and extracted the corresponding features at the centre of those clusters – the list of the most informative features is given in **Table S12** and the magnitude probability density plots of the central features with the highest probability are shown in **Fig. 5f**. We found that the classification was essentially dominated by two timeseries features, i.e., the ‘information gain’ feature, which estimates how easy it is to predict a data point in the time series from the preceding data points, and the ‘quadratic fit of power spectrum cumulative sum’ feature, which characterizes the power spectrum of the time series. The increase in ‘quadratic fit of power spectrum cumulative sum’ during stimulation can be simply attributed to the drop in the spectral peak at the tremor’s frequency. In contrast, the increase in ‘information gain’ during stimulation revealed a loss of linear dependency between consecutive datapoints of the tremor movement, i.e., a loss of temporal coherence.

To specifically test whether the stimulation-induced change in the tremor amplitude was associated with a change in temporal coherence, we computed the change in the magnitude-squared coherence during the stimulation period relative to the baseline period in the ‘decrease’ and the ‘increase’ datasets as well as in a dataset consisting of all the trials with sham stimulation (‘sham’). We found that the temporal coherence in the tremor frequency-band decreased in the ‘decrease’ dataset and increased in the ‘increase’ dataset during the stimulation, however, it did not change in the ‘sham’ dataset (**Fig. 5g**). The change in the tremor amplitude in the ‘decrease’ dataset, but not in the ‘increase’ dataset, was correlated with the change in the tremor temporal coherence. The change in the tremor amplitude in the ‘sham’ dataset was also positively correlated with the change in the tremor temporal coherence, however, with a smaller slope of the linear regression (**Fig. 5h**). The decrease in temporal coherence in the ‘decrease’ dataset was correlated with the onset of the stimulation and was maintained during the duration of the stimulation (**Fig. 5i**).

### Possible mechanism of action via perturbation of spiking regularity in the olivocerebellar loop

To explore the neurophysiological mechanism of our stimulation strategy we used a computational model of the CCTC network under ET condition^15^ and tested the effect of phase-locked stimulation of the cerebellum on the spiking dynamics. **Fig. 6a** shows a schematic representation of the model. As in our previous study^15^, the power spectrum density (PSD) between 4 –12Hz of the ventral intermediate thalamus (Vim) was used as a proxy to the tremor activity. **Fig. 6b** shows a representative PSD of the Vim under normal and ET conditions with a pathologic oscillation peak at 7Hz.

**Fig. 6.**
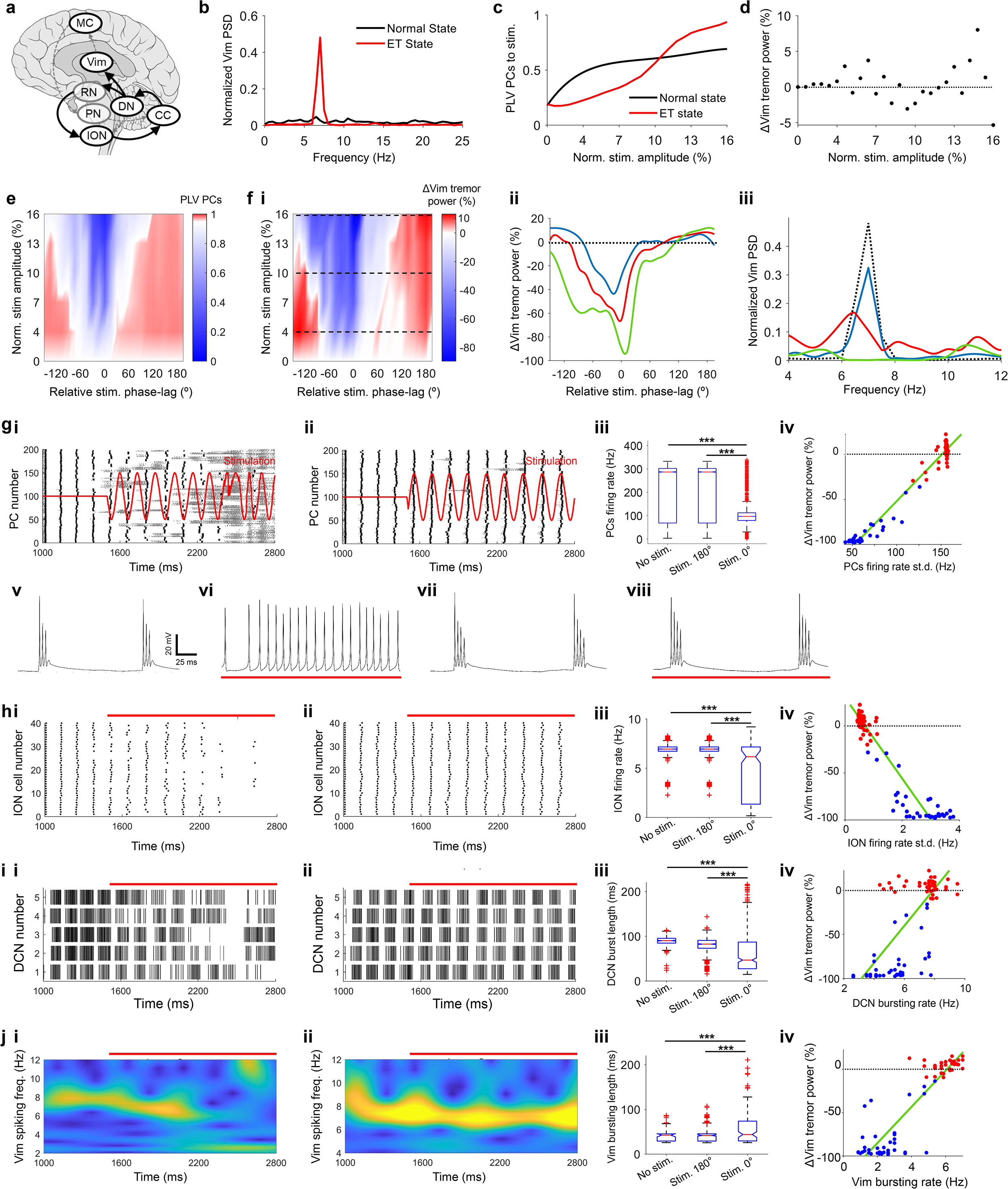
Mechanism of phase-locked cerebellar stimulation in ET, probed via neurophysiological modelling. Cortico-cerebello-thalamo-cortical (CCTC) network model was simulated under ET condition^15^. Sinusoidal currents were applied to the modelled Purkinje cells to simulate transcranial electrical stimulation of the cerebellum. In case of phase-locked stimulation, the instantaneous phase of the sinusoidal currents was adjusted online to achieve a fixed phase-lag to the instantaneous phase of the spike count trace of the thalamocortical neurons in the ventral intermediate thalamus (bandpass filtered at 6-10 Hz), computed via ecHT. **a**, Schematic of the CCTC network model. CC, cerebellar cortex containing Purkinje cells (PC) and granular layer cells; DN, dentate nucleus containing deep cerebellar neurons (DCN) and the nucleo-olivary (NO) neurons; ION, inferior olive nucleus; RN, red nucleus; PN: pontine nucleus; Vim, ventral intermediate nucleus of the thalamus including thalamocortical (TC) neurons; MC, motor cortex containing pyramidal neurons (PYN) and fast-spiking interneurons (FSI); solid arrows, pathways associated with the generation of ET; grey dashed arrows, pathways not associated with the generation of ET. See Zhang et al.^15^ for a detailed description of the model. **b**, Normalized power spectral density (PSD) of the spike count trace of TC neurons in the Vim during normal tremor free condition, black line, and ET condition, red line (tremor frequency 7Hz). ET condition is used throughout the rest of the figure. PSD values were normalized to total power between 0 Hz and 25 Hz. Simulation duration, 11.5 s. **c-d**, Sinusoidal current stimulation of PCs without phase-locking. Stimulation frequency, 7Hz. Each current amplitude value was simulated for 10 s preceded by 1.5 s stimulation-free period. Current amplitude values were normalized to the amplitude of the endogenous synaptic current measured at rest under ET state over 4000 ms. **c**, Modulation of PCs phase-locking value (PLV) versus normalized amplitude of stimulation (see **Online Methods** for details of PLV computation) during normal condition, black line, and ET condition, red line. **d**, Modulation of tremor PSD in Vim (i.e., maximal PSD between 4 −12 Hz), showing percentage change in tremor PSD with respect to ET without stimulation versus normalized amplitude of stimulation. **e-k**, Sinusoidal current stimulation of PCs phase-locked to Vim’s tremor spike train. Stimulation as in (c-d) but instantaneous phase was adjusted online to maintain a fixed phase-lag of 0° to 330° (30° phase increments; see **Online Methods** for details of online phase-locking computation). Phase lag values are expressed relative to 330° phase lag showing the largest reduction in tremor amplitude and wrap to ±180°. **e**, Modulation of PCs PLV, showing colormap of PLVs versus normalized amplitude and phase of stimulation. **f**, Modulation of tremor PSD in Vim, shown are (i) colormap of percentage change in tremor PSD versus normalized amplitude and phase of stimulation; black dashed line, normalized stimulation amplitude of 4%, 10% and 16%, (ii) line plots of percentage change in tremor PSD in Vim versus stimulation phase for amplitudes 4% (blue line), 10% (red line), and 16% (green line), and (iii) Normalized Vim PSD during stimulation with a relative phase of 0° and normalized amplitudes of 4% (blue line), 10% (red line), and 16% (green line); black dashed line, without stimulation. **g**, Representative spiking activity of PCs, showing (i) spike raster plot during stimulation at 16% normalized amplitude and 0° relative phase-lag (black) and waveform of stimulating current (red); (ii) same as (i) but at 180° relative phase-lag; (iii) statistics of PCs’ instantaneous spiking rate (i.e., 1/inter-spike-interval) during no stimulation (‘No stim.’), stimulation at 180° relative phase-lag (‘Stim. 180°’), and stimulation at 0° relative phase-lag (‘Stim. 0°’); red line, median; box edges, 25% and 75% percentiles; *** p<0.0001; significance was assessed using Bartlett’s test and post-hoc two-sample F test with Bonferroni corrections for multiple comparisons; stimulation started after 1,500ms and lasted till the end of the simulation; (iv) correlation between change in Vim tremor PSD and standard deviation (st.d.) of PCs’ instantaneous spike firing rate; c=-154, m=1.3, R^2^ =0.97 (c, line y-intercept; m, line slope; green line, linear fit); stimulation at 0° relative phase-lag resulted in a breakdown of complex spikes (high median instantaneous spiking rate due to short inter spikelet intervals, high st.d. instantaneous spiking rate due to long inter burst intervals) and restoration of regular simple spikes; (v-vi) representative spike train of a PC before (v) and during (vi) stimulation at 0° relative phase-lag; (vii-viii) same as (v-vi) but for 180° relative phase-lag. **h**, same as (g) but showing results for ION neurons; red horizontal bar indicates stimulation period; c=30, m=-43, R^2^ =0.9. **i**, same as (h) but showing results for DCNs, (iii) change in DCNs’ spiking temporal coherence z-scored relative to mean and st.d. during baseline (i.e., pre-stimulation period), and (iv) correlation between change in Vim tremor PSD and change in DCNs’ temporal coherence z-scored as in (iii); c=-5.8, m=14, R^2^ =0.97. **j**, same as (i) but showing results for TC neurons in Vim, (i-ii) representative spiking spectrograms during the same stimulation conditions as in g-i(i-ii); (iii-iv); c=2.5, m=25, R^2^ =0.95.

We first assessed the effect of periodic electrical stimulation of the cerebellum without phase-locking by adding sinusoidal input currents to the Purkinje cells (PCs) which were chosen as the stimulation target due to their abundance in the cerebellar cortex^6^ and their high excitability (see **Fig. S2a-b** for comparison with granular cells). We quantified the change in the spiking activity due to stimulation at the tremor frequency and a range of amplitudes evoking small sub-threshold depolarization as expected in our experiment (see **Online Methods** for details). We found that the stimulation entrained the PCs with an efficiency that increased with the current amplitude (**Fig. 6c**). However, the entrainment of the PCs resulted in only a small decrease (<5%) or increase (<10%) in the tremor PSD of the Vim (**Fig. 6d**, see also **Fig. S2c** for representative spiking activity).

We then assessed the effect of phase-locking the stimulation to the tremor activity by computing the instantaneous phase of the TC neurons’ spike-train online using ecHT and adjusting the phase of the sinusoidal currents to maintain a fix phase-lag as we did in our experiment (see **Online Methods** for details). We found that in this case, a narrow range of stimulating phase-lag values were capable of perturbing the synchronous activity of the PCs (**Fig. 6e**) resulting in a large decrease (up to ~95%) in the tremor PSD of the Vim (**Fig. 6f**). Higher current amplitudes resulted in a larger decrease in tremor-PSD and a wider range of efficacious phase-lags. Stimulating phase-lag values outside this range could result in a small increase (up to ~10%) in the tremor PSD of the Vim. Increasing the number of cells in the model affected the range of efficacious phase-lags but not the resulted drop in tremor PSD (**Fig. S2d**).

At efficacious phase-lags, the hyperpolarizing phase of the stimulating currents was consistently aligned with the onset time of the complex spikes in the PCs, resulting in a gradual suppression of the periodic complex spiking and restoration of more physiological simple spiking (**Fig. 6g**). The perturbation of the complex spiking in the PCs disrupted the periodicity of the spiking in the inferior olivary nucleus (ION) neurons (**Fig. 6h**) and the temporal coherence of the bursting activity in the deep cerebellar neurons (DCNs; **Fig. 6i**) thus ceasing the tremulous coherent drive to the Vim of the thalamocortical loop (**Fig. 6j**). In contrast, at 180° relative phase-lags, the depolarizing phase of the stimulating currents was consistently aligned with the onset time of the complex spikes in the PCs, resulting in a small augmentation of the periodic complex spiking that led to a small reinforcement of the aberrant tremulous drive to the Vim of the thalamocortical loop (**Fig. 6g-j**).

## Discussion

In this paper we presented the ecHT strategy to compute the instantaneous phase of oscillatory signals in real-time and validated it using both simulation and measurements with pathologic oscillatory brain activity, i.e., ET. The ecHT strategy is based on the application of a causal bandpass filter to the DFT of the analytic signal to mitigate the distortion, known as the Gibbs phenomenon, from its end. Given the widespread use of the Hilbert transform to compute the instantaneous attributes of oscillatory signals^10^, the possibility for real-time computation using ecHT opens exciting opportunities. Future studies may be able to improve the accuracy of the ecHT by adjusting, online, the central frequency of the bandpass filter to the instantaneous frequency of the signal, computed e.g., via a time derivative of the instantaneous phase.

Other frequency-domain and time-domain filters have been previously proposed to mitigate the Gibbs phenomenon from finite signals with a discontinuity^16^ but these filters restore the DFT only away from the discontinuity itself^17^. There have also been reports of restoring the endpoint of the analytic signal using recursive models, such as autoregression^18^ or polynomial fitting^19^ however their high runtime complexity has limited their use in applications that require real-time computation using conventional digital hardware.

We then reported an early-stage proof of concept of non-invasive neuromodulatory strategy to ameliorate ET via transcranial cerebellar stimulation that is phase-locked to the endogenous tremor movement computed in real-time using ecHT. The range of phases that were efficacious in suppressing the tremor in our stimulation was small but varied between patients and within patients between days of experiments perhaps due to differences in the electrode-skin capacitance. Our results exemplify the importance of accurate phase-locking to successfully induce a reduction in tremor amplitude. The fact that the tremor amplitude continued to drop during the stimulation period suggests that a longer stimulation period may yield an even larger suppression. The sustained drop in tremor amplitude after the end of the stimulation period may hold potential for a therapeutic effect via neural plasticity.

Given the large proportion of ET patients that discontinue oral medication due to insufficient benefits and/or adverse effects^6^, and the risks associated with invasive lesioning and DBS therapies, the prospect for non-invasive brain stimulation treatment is very enticing. Future studies with larger patient cohorts and longer stimulation periods, are needed to better pinpoint the magnitude and duration of the tremor reduction and to assess the safety profile. There has been an original report that showed that phase-locked neuromodulation of the motor cortex can ameliorate tremor in Parkinson’s disease (PD) patients^20^. Although both ET and PD are caused by aberrant oscillations in the motor system, their anatomical origins and degree of coupling between the central oscillators are very distinct^21^. There has been evidence for cellular and molecular changes in DCN and PC in ET patients^22^. Our results emphasize the critical role of the DCN as the main cerebellar efferent in both the generation and the maintenance of ET oscillations and demonstrates its sensitivity to modulation via PC activity. There has also been a report that a periodic stimulation of the motor cortex at the tremor frequency without phase-locking, can entrain the phase of ET in patients undergoing DBS with an efficiency that was correlated to the somatosensory sensation underneath the electrodes^23^. In our study, the changes in the circular phase distribution and amplitude of the tremor were not dependent on the subjective sensation of the patients.

The mechanism of action of our neuromodulatory strategy, as revealed via feature-based statistical learning and neurophysiological modelling, involves direct perturbation of the cascade of coherent activities that generate the tremor oscillation in the olivocerebellar loop. This mechanism of action differs from existing Vim DBS therapy for ET which masks the tremor oscillation in the thalamocortical loop but does not mitigate its generation in the olivocerebellar loop^15^. In the future, neuromodulatory strategies that target the temporal coherence of the pathology may offer new opportunities to treat a wide range of brain disorders underpinned by aberrant synchronous oscillations.

## Online Methods

### Endpoint corrected Hilbert transform (ecHT)

A discrete analytic signal is most accurately and efficiently computed by deriving the discrete Fourier transform (DFT) of the signal, zeroing the Fourier components of the negative frequencies and doubling the ones of the positive frequencies, and constructing the analytic signal using the inverse discrete Fourier transform (IDFT)^11^. However, Gibbs phenomenon distortion^9^ in the derivation of the analytic signal at the ends of finite-length signals has rendered an accurate computation of the instantaneous phase and envelope amplitude at the last data point impossible^12^. Since the Gibbs phenomenon stems from a nonuniform convergence of the DFT at a discontinuity between the beginning and the end of the analytic signal^24^, we hypothesized that by applying a causal bandpass filter to the DFT of the analytic signal we would establish a continuity between the two ends of the signal and remove the distortion selectively from the end part of the signal. The bandpass feature of the filter reduces extraneous DFT coefficients, limiting the oscillatory properties to the target frequency-band, while balancing the phase-lag introduced by the low-pass component of the filter with the phase-lead introduced by the high-pass component of the filter. The causality feature of the filter restores the linear increment of the phase at the end of the analytic signal by projecting the oscillatory properties from the adjacent, non-distorted data points. Since the DFT treats finite sampled signals as if they were replicated periodically, the projection of the oscillatory properties would continue through the beginning of the signal, thus forcing a continued increment of the phase from the restored signal end to its beginning. The runtime complexity of the filtering is *O*(*n/2*), where *n* is the number of frequency points, is lower than of the *O*(*n · log*(*n*)) fast Fourier transform (FFT) and inverse fast Fourier transform (IFFT) that dominates the computation of the analytical signal.

### Simulation of ecHT

Simulation of ecHT was done in MATLAB (MathWorks Inc). A discrete oscillatory test signal *y_i_*[*n*] = *A_i_cos*(2*πf_i_n* − Ø_*i*_) was generated (*i*. being the signal number) over a finite time interval, where 0 > *n* > *N* − 1 was the time point number and *N* was the total number of time samples, *A_i_* was the envelope amplitude of the signal, *f_i_* was the frequency of the signal, and Ø_*i*_ was the phase delay of the signal.

The analytic signal was computed by first computing the Fourier representation *y_i_*[*k*] of the signal using MATLAB’s fast FFT function (‘fft’), where 0 > *k* > *K* − 1 was the frequency bin number and *K* was the total number of frequency samples. Then, generating the Fourier representation *Z_i_*[*k*] of the analytic signal by zeroing the Fourier components of the negative frequencies and doubling the Fourier components of the positive frequencies, i.e.,

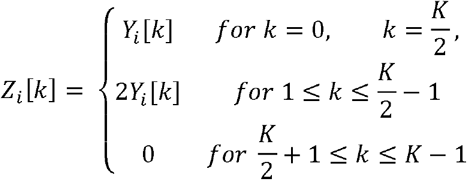

If ecHT was applied, the Fourier representation of the analytic signal *Z_i_*[*k*] was multiplied with the response function σ[*k*] of a Butterworth bandpass filter that was obtained using MATLAB’s frequency response of digital filter function (‘freqz’) from the filter’s impulse response coefficients generated using MATLAB’s Butterworth filter design function (‘butter’). Finally, the analytic signal *z_i_*[*n*] was computed from its Fourier representation *Z_i_*[*k*] using MATLAB’s IFFT function (‘ifft’). The phase of the signal at the last data point was computed via 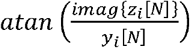, where *imag*{*Z_i_*[*N*]} is the imaginary part of the analytic signal, i.e., the Hilbert transform of the original signal, and was compared to the actual phase of the signal at the last data point, i.e., 2*πf_i_N* − Ø_*i*_. The amplitude of the signal at the last data point was computed via 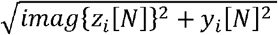 and was compared to the actual amplitude of the signal at the last data point, i.e., *A_i_*.

### Feasibility study of cerebellar electrical stimulation phase-locked to ET

The study design included a cross-over, double-blinded randomization of conditions with a sham and an active control. It was approved by the local research ethics committee in accordance with the declaration of Helsinki. All participants provided written informed consent prior to study participation.

#### Participants

Eleven ET patients (3 females) were recruited from the outpatient department of the UK National Hospital of Neurology and Neurosurgery, London. All patients fulfilled the diagnostic criteria for ET according to the Tremor Investigation Group and consensus statement of the Movement Disorder Society^25^ and were on a stable treatment regime for their tremor for at least 30 days prior to the experiment. See **Table S1** for demographic and clinical information. Experiments were performed after overnight withdrawal of tremor medication during a single study visit in the dominant hand, or in case of slight asymmetry in the hand with the larger tremor amplitude. There were no drop-outs or adverse events noted.

#### Participants (second cohort)

Seven ET patients (4 females) were recruited as in the original to test whether their response can be predicted via the feature-based approach developed in the original study. See **Table S9** for demographic and clinical information. Experiments were performed as in the original cohort.

#### Experiment design

The experiment consisted of eight stimulation conditions, i.e., six sinusoidal stimulating currents that are phase-locked to the tremor movement at different phase lags (i.e., 0°, 60°, 120°, 180°, 240° and 300°), a control sinusoidal current at the tremor frequency but without phase-locking, and a sham stimulation condition. Each stimulation condition was applied in a block (i.e., trial) of 60s during which the patients sat in an armchair and were instructed to maintain a tremor evoking posture, i.e., stretched, elevated arm with fingers parted, while their tremor movement was measured (see details below). The 60s block included a 15s of a baseline period, a 30s of a stimulation period (including 5s of ramp-up and 5s of ramp-down at the beginning and end of the stimulation, respectively) and a 15s of post-stimulation period. In sham stimulation blocks, the current was set to zero after the 5s of ramp-up. Each 60s block was preceded by a short (~4s, 2048 data samples) calibration recording also in a tremor evoking posture to compute the tremor frequency and amplitude at the onset of the block (see details below). The eight stimulation conditions were applied consecutively with a 30s rest interval between conditions. The sequence of eight stimulation conditions was repeated four times (apart from one patient in which they were applied three times due to fatigue) in a random order with 10min rest period between sequences. The rest interval between conditions and the rest period between sequences were occasionally extended slightly if the patients requested.

#### Measurement and real-time computation of instantaneous tremor phase via ecHT

Tremor movements were measured using a 3-axis analog microelectromechanical system (MEMs) accelerometer (MMA7361, Freescale Semiconductor, Inc.; operated at a sensitivity range of ±1.5G) that was attached to the proximal phalangeal segment of the middle finger using a custom-made adapter. The 3-axis acceleration measurements were sampled using three analog-to-digital converters (ADCs) of a microcontroller (Arduino Due with an Atmel AT91SAM3X8E processor and a single ARM Cortex M3 core; operated at a clock rate of 84 MHz) at a rate of ~500Hz and an amplitude resolution of 12-bit, and the vector amplitude sum of the three axes was computed and stored in a running window of 128 samples. The instantaneous phase and amplitude of the tremor movement, i.e., at the last sample of the running window, were computed in real-time and at the same rate, using ecHT that was implemented on the microcontroller. The ecHT implementation had a 2nd order Butterworth bandpass filter (2nd order low pass, 2nd order high pass) with a bandwidth that was equal to half the frequency of the tremor and was centered at the frequency of the tremor. The frequency of the tremor was computed using FFT from a short calibration measurement of 2048 samples (i.e., frequency resolution of ~0.25Hz) before each 60s stimulation block. The sampled tremor movement measurement was logged to a laptop, together with the ecHT setting and the tremor frequency and amplitude computed during calibration, using a Processing script that was also used to interface with the microcontroller.

#### Transcranial stimulation of ipsilateral cerebellum

Sinusoidal stimulating currents were generated by first producing voltage waveforms, pseudo-differentially via two digital-to-analog converters (DACs) of the microcontroller (with an amplitude range of ±1V and an amplitude resolution of 12-bit) and then feeding them to an isolated bi-phasic current source (DS4, Digitimer Ltd; operated at an input range of ±1V and an output range of ±1mA or ±10mA). The frequency of each voltage waveform was equal to the frequency of the tremor computed before each 60 s stimulation block as mentioned above. To phase-lock a stimulating current to the ongoing tremor movement, the phase of the voltage waveform was adjusted, at the same rate of 500Hz, to maintain a fixed phase lag to the computed phase of the last acceleration sample. The amplitude of the stimulating currents was 2.7 ±1 mA (mean ±st.d.) across the patient cohort, (the amplitude was individually adjusted for each patient below any discomfort level due to extraneous somatosensory stimulation underneath the `electrodes`). To reduce risk of extraneous high-frequency stimulation due to low signal-to-noise (SNR) level, the amplitude of the voltage waveform was set to zero when the amplitude of the last acceleration sample was <1% of the amplitude during the short calibration measurement before each 60s stimulation block. The generated stimulating voltage waveforms were logged to a laptop together with the tremor movement measurements using the same Processing script.

The stimulating currents were applied transcranially to the ipsilateral cerebellum transcranially via a 2 x 2 cm^2^ skin electrode (Santamedical, 2” X 2” carbon electrode pad with Tyco gel that was cut to the specified dimensions) that was placed 10% nasion-inion distance lateral to inion (i.e., above the cerebellar lobule VIII) and was paired with a 5.08 x 5.08 cm^2^ skin electrode (the same carbon electrode pad but was not cut) that was placed over the contralateral frontal cortex between F3-F7 or F4-F8 of the international 10-20 system. Before the placement of the electrodes, the scalp skin was prepared using 80% Isopropyl alcohol and an abrasive skin gel (NuPrep, Weaver and Company Inc), and a conductive paste (Ten20, Weaver and Company Inc) and/or a conductive gel (CG04 Saline base Signa gel, Parker Laboratories Inc) was deposited at the target locations. The resistance between the electrodes was maintained below 8 kOhm.

### Analysis of stimulation phase lag

Analysis of the stimulation phase lag was done in MATLAB. The tremor movement trace of each 60 s block was filtered with the same filter settings that were used in the real-time computation, i.e., a 2nd order Butterworth bandpass filter with a bandwidth that was equal to half the frequency of the tremor and centered at the frequency of the tremor computed and logged at the short calibration period preceding each block. The instantaneous phase of the stimulating waveform trace and the instantaneous phase of the filtered tremor movement trace were computed using MATLAB’s ‘hilbert’ function, and the instantaneous phase lag between the two traces was calculated and then epoched in intervals of 1s. The stimulating trace in the sham condition was a virtual sinusoidal waveform at the tremor frequency.

The statistics and statistical tests of the phase lag values were computed, using MATLAB’s CircStat toolbox^13^, in the following periods – the whole stimulation period (20s since 5s ramp-up time and the 5s ramp-down time at the beginning and the end were excluded, respectively), the first half of the stimulation period (10s since 5s ramp-up time was excluded), the second half of the stimulation period (10s since 5s ramp-down time was excluded). First, the unimodality of the phase distribution of each stimulation condition was validated using Watson’s test against a von Mises distribution (set phase 0°, p<10^−5^; 60°, p<10^−5^; 120°, p<10^−5^; 180°, p<10^−5^; 240°, p<10^−5^; 300°, p<10^−5^; no phase-lock, p=0.6). The phase distribution during stimulation with phase-locking was not different from von Mises distribution but since the phase distribution during stimulation without phase-locking was different from von Mises distribution, we used non-parametric statistical tests. Next, the circular spread of the phase distribution of each stimulation condition was quantified by computing the length of the mean resultant vector *R* and its uniformity was assessed using the Omnibus test. Then, the difference between the mean phase of the stimulation conditions was assessed using Fisher test and the difference between the mean resultant vector length *R* of the stimulation conditions was assessed using ANOVA with post-hoc analysis using Wilcoxon signed-rank test. Finally, the effect of the tremor parameters, i.e., amplitude and frequency, on the length of the mean resultant vector *R* was assessed via Pearson correlation.

### Analysis of change in tremor amplitude

Analysis of the tremor amplitude was done in MATLAB. The tremor trace of each 60s block was filtered as in the ‘Analysis of stimulation phase lag’. The instantaneous amplitude was computed using MATLAB’s ‘hilbert’ function and was epoched in intervals of 1s. To express the tremor amplitude relative to the amplitude of the baseline period, the amplitude value of each epoch was z-scored by subtracting the mean value during the baseline period and then dividing by the st.d. of the value during the baseline period.

The statistics and statistical tests of the tremor amplitude values were computed in the following periods – the baseline period (10s between 3s and 13s from block onset), the whole stimulation period (as in ‘Analysis of the stimulation phase lag’), the first half of the stimulation period (as in ‘Analysis of the stimulation phase lag’), the second half of the stimulation period (as in ‘Analysis of the stimulation phase lag’), and the post-stimulation period (10s between 3s and 13s from stimulation offset). To assess the change in the tremor amplitude relative to the change in the tremor amplitude during the sham stimulation condition, the z-score amplitude values during stimulation and during post-stimulation periods of each stimulation condition were subtracted by the corresponding median z-score values of the sham stimulation condition.

To assess the effect of phase-locking the stimulation to the tremor movement, the change in the tremor amplitude due to stimulation with phase-locking and without phase-locking was analysed. First, the change in the tremor amplitude due to each type of stimulation, i.e., without phase-locking and with phase-locking (data from all six phase-lags of stimulation was combined) was assessed across the patient cohort in each epoch using unpaired t-test. Next, the change in tremor amplitude of individual patient due to each stimulation condition was assessed (i.e., data including four repetition trials from each phase-lag of stimulation was treated separately) during stimulation and post-stimulation periods using unpaired t-test as well as using surrogate distributions (i.e., 1000 z-scores values with the same st.d. but zero mean value), where the p-value threshold of the stimulation conditions with phaselocking (but not without phase-locking) were Bonferroni corrected for the six phase lag conditions. Then, the number of patients that showed statistically significant increase/ decrease of z-score amplitude was assessed using Fisher’s exact test against the number of patients who did not show a change in the z-score tremor amplitude (patients could have a significant increase of z-score in one phase-lag and a significant decrease of z-score in another phase-lag). Finally, the z-score amplitude of the sub-group of subjects that showed a statistically significant increase/decrease of z-score amplitude was assessed using unpaired t-test.

To assess the effect of the phase lag value during stimulation, the change in the tremor amplitude due to stimulation with different phase lags was analysed. First, the change in the tremor amplitude due to each phase-lag of stimulation was assessed across the patient cohort during the stimulation period using unpaired t-test. Next, the change in tremor amplitude of individual patient was assessed during stimulation again using unpaired t-test. Then, the number of patients that showed a statistically significant increase/ decrease of z-score amplitude was assessed using Fisher’s exact test. Finally, to account for differences in phase response across patients, the phase lags were expressed relative to the phase lag that resulted in the largest reduction in the tremor amplitude, and the change in tremor amplitude of individual patient and the number of patients with statistically significant change were reanalysed.

### Prediction of patients’ response to stimulation from features of tremor movement

#### Dataset

Time-series of tremor movement during the baseline period, i.e., 10s (5000 data points) from 5s after the onset of tremor posture till 5s before the onset of the phase-locked stimulation, were extracted from all the recorded trials with phase-locked stimulation, resulting in a dataset of 301 time-series trials (28 trials per patients except patient 3 in which only 21 time-series trials were recorded). The time-series were assigned a ‘responder’ or a ‘non-responder’ label if the patient responded or did not respond to the stimulation, respectively. A patient was conservatively labelled as a ‘responder’ if his/her tremor amplitude decreased in at least one of the tested stimulation phases relative to sham and did not increase in any of the tested stimulation phases relative to sham, and was labelled a ‘non-responders’ if his/her tremor amplitude increased in at least one of the tested stimulation phases relative to sham or did not change in any of the tested stimulation phases relative to sham.

#### Extraction of time-series features

For each time-series trace, 7873 features were computed using the highly comparative time-series analysis (*hctsa*) ^14^, resulting in a 301 x 7873 feature matrix. The computed features included autocorrelations, power spectra, wavelet decompositions, distributions, time-series models (e.g. Gaussian Processes, Hidden Markov model, autoregressive models), information-theoretic quantities (e.g. Sample Entropy, permutation entropy), non-linear measures (e.g. fractal scaling properties, nonlinear prediction errors) etc. All features with infinity or not a number (NaN) values and features with zero variance across the dataset were removed from the feature matrix, resulting in a reduced feature matrix of 301 x 6196. The value of each feature was individually normalized to the interval [0,1].

#### Classification

The feature space was partitioned, i.e., classified, using a linear Support Vector Machine (SVM) classifier, implemented with the *classify* function of MATLAB’s *Statistics Toolbox*, which returned a threshold that optimally separated the two classes, i.e., ‘responders’ and ‘non-responders’ time-series. The accuracy of the classification was quantified by first computing the balanced classification accuracy 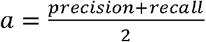, and then computing the harmonic mean of precision and recall, i.e., *F*_1_ score, 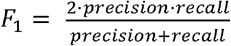, where precision is the fraction of true positive classified samples over the total of positively classified samples and recall is the fraction of true positive classified samples over the total true positive and false negative classified samples. The classification was performed using a 10-fold cross-validation to reduce bias and variance.

#### Performance-based feature selection

The univariate classification performance of each feature was evaluated against the class labels. A subset of 40 features with the highest single-feature classification accuracy was selected. To reduce the redundancy within the subset of features, the Pearson correlation distance, *d_ij_* = 1 - *ρ_ij_* was computed for each pair of features, where *ρ_ij_* is the Pearson correlation coefficient between feature *i*. and feature *j*, and a hierarchical clustering was performed using a complete linkage threshold of 0.2, resulting in clusters of features that were inter-correlated by *ρ_ij_* > 0.8. The clusters of highly correlated features were then represented by the feature that was located most centrally within the cluster (i.e., at the cluster’s centre).

#### Feature-based prediction of patient response

The centroid of individual patients in the feature space (including the extracted 14 most informative features) was computed by averaging the feature values across the corresponding trials. The centroid of the patient class (i.e., ‘responders’ or ‘non-responders’) in the same feature space was computed by averaging the features values across the corresponding trial dataset. The Euclidean distance between feature centroids was computed with *pdist* function of MATLAB.

#### Visualization using principal component analysis

To facilitate visualisation of the feature space, principal component analysis (PCA) was performed. In this case, a covariance matrix was computed for the normalized set of features from which the eigenvectors and eigenvalues were extracted. Each principal component was constructed as a linear combination of the initial features. The first two principal components were then used to display 2D scatter plots of the features.

### Change in features of tremor movement due to stimulation

#### Dataset

Time-series of tremor movement during stimulation (10s; 5000 data points; from 10s after the onset of stimulation till 10s before the offset of stimulation) and during baseline (10s; 5000 data points; same as in ‘Classification and prediction of patients’ response to stimulation’) from all trials with phase-locked stimulation (301 traces of stimulation and baseline each) were extracted and assigned a ‘stimulation’ class label or a ‘baseline’ class label, respectively.

The ‘stimulation’ and ‘baseline’ time-series were then divided into three datasets according to the change in the tremor amplitude during stimulation, i.e., ‘decrease’, traces in which the tremor amplitude decreased during stimulation relative to sham (58 time-series of stimulation and baseline each, 11 subjects); ‘increase’, time-series in which the tremor amplitude increased during stimulation relative to sham (51 time-series of stimulation and baseline each, 10 subjects); ‘no-change’, traces in which the tremor amplitude did not change during stimulation relative to sham (192 time-series of stimulation and baseline each, 11 subjects). In addition, in a subset of the analysis, the same ‘stimulation’ and ‘baseline’ tremor traces were extracted from all the blocks with sham stimulation (‘sham’; 43 time-series of stimulation and baseline each, 11 subjects).

#### Extraction of time-series features, classification, and performance-based feature selection

Same as in ‘Classification and prediction of patients’ response to stimulation’.

#### Temporal coherence analysis

The tremor temporal coherence versus frequency of each tremor trace was quantified by computing the magnitude squared coherence across 1s epochs during ‘stimulation’ period and ‘baseline’ period using MATLAB’s *mscohere* function with a frequency range of 0 to 31 Hz and a 1Hz frequency resolution. The computed values during ‘stimulation’ were then z-scored relative to the mean and st.d. of the values during ‘baseline’. The tremor temporal coherence at the tremor frequency band was quantified by computing the mean z-score across the 4 – 8 Hz frequency bins. The tremor temporal coherence versus time of each tremor trace was quantified by computing the magnitude squared coherence between 1s epoch and its preceding one during ‘stimulation’ period and ‘baseline’ period using the same MATLAB’s *mscohere* function, z-score the ‘stimulation’ values relative to ‘baseline’ in the same way, and then computing the mean z-score across the 4 – 8 Hz frequency bins. Statistical significance of magnitude squared coherence at a frequency bin was characterized for each dataset (i.e., decrease’, ‘increase’, and ‘sham’) using unpaired t-test with Bonferroni corrections for multiple comparisons of frequency bins and datasets.

### Neurophysiological modelling

CCTC network model under ET condition was simulated as in Zhang et al.^15^. The model is available on ModelDB (http://modeldb.yale.edu/257028). It consisted of 425 single-compartment, biophysics-based neurons from the olivocerebellar and thalamocortical loops, including 40 inferior olivary nucleus (ION) neurons in the brainstem, 200 Purkinje cells (PCs) and 20 granular layer clusters (GrL) in the cerebellar cortex, 5 glutamatergic deep cerebellar projection neurons (DCNs) and 5 nucleoolivary (NO) neuron in the dentate nucleus, 5 ventral intermediate thalamus (Vim) thalamocortical (TC) neurons, 100 pyramidal (PYN) neurons, and 10 fast-spiking interneurons (FSI). As in our previous study^15^, the ET condition was simulated by reducing the conductivity and increasing the decay time of the PCs’ GABAergic currents to the DCN, which mimics the loss of GABAA α_1_-receptor subunits and an upregulation of α_2_/α_3_-receptor subunits in the cerebellum. Five instances of the model were considered and for each instance, simulations were repeated under normal condition, ET condition with no stimulation, and ET condition with stimulation of the cerebellum. Each simulation lasted 11,500ms (integration step, 0.0125ms). ET condition was initiated after 1000ms and stimulation started after 1,500ms and lasted till the end of the simulation.

To simulate the cerebellar stimulation, a current *I_stim_* was added to all the PCs in the model. *I_stim_* was sinusoidal with a frequency that is equal to the frequency of the ET and amplitudes between 1-5pA evoking small subthreshold depolarizations expected in our experiment. Specifically, *I_stim_* with an amplitude of 1pA induced a periodic depolarization of ~0.5mV amplitude in the single-compartment PC model that was similar to the depolarization that was induced by an extracellular electric field with an amplitude of 2V/m, predicted from our FEM modelling of the experiment (**Fig. 2b**), in the multi-compartment PC model (**Fig. S2a-b**). The amplitude of *I_stim_* was normalized to the average amplitude of the endogenous synaptic current to PCs, measured under ET state over 4000ms (see also Perkel et al.^26^), with *I_stim_* of 1pA equals 4% of the average endogenous synaptic current to PCs. To phase lock the sinusoidal current to the ET oscillation, first the spike count trace of the TC neurons of the Vim was computed with a temporal resolution of 1ms and then filtered using a 2nd order Butterworth bandpass filter with cut-off frequencies of 6Hz and 10Hz). Then, the instantaneous phase of the spike count trace was computed online every 10ms using ecHT on a running window of 1000ms, and the phase of the stimulating current was adjusted at that time points to maintain the target phase lag.

#### Computation of PCs phase-locking value

First, the spike count trace of the PCs was computed with a temporal resolution of 1ms (spikes were summed across PCs) and low pass filtered using a 2nd order Butterworth filter with a cut-off frequency of 30 Hz. Then, the instantaneous phases of the spike count trace and the stimulating current were computed offline using MATLAB’s ‘hilbert’ function, and the instantaneous phase lag between the two was calculated every 1ms. The phase-locking value (PLV) of each PC was computed as in Lachaux et al.^27^ and then averaged across the PCs.

#### Computation of Vim power spectrum density

First, the spike count trace of the TC neurons in the Vim was computed with a temporal resolution of 1ms (spikes were summed across TC neurons). Then, the power spectral density (PSD) of the spike count trace was computed using Welch’s method with 2,000ms Hanning window and 1,000ms overlap, and normalized to the total power between 0Hz and 25Hz. Tremor PSD was estimated as the peak PSD at the tremor frequency band, i.e., between 4 −12 Hz.

#### Computation of DCN and Vim temporal coherence

The spike trains of the DCN and TC neurons of the Vim were low pass filtered using a 2nd order Butterworth filter with a cut-off frequency of 30 Hz, and the magnitudes squared coherence were computed using MATLAB’s *mscohere* function with a frequency range of 0 to 30 Hz. Then the magnitude squared coherence in DCN and Vim during stimulation was expressed relative to baseline by subtracting the mean value during baseline and dividing by the st.d. value during baseline, i.e., z-score.

### Transcranial electric field modelling

Finite element method (FEM) electromagnetic simulations were performed in Sim4Life V.4 (ZMT ZurichMedTech AG, Zurich), using a quasi-static ohmic-current solver. Electrodes were created within the platform using Sim4Life’s CAD functionalities and applied to the scalp of the MIDA anatomical head model^28^. Dirichlet (voltage) boundary conditions were assigned to the electrodes, and tissues electrical conductivities were assigned according to the IT’IS LF database^29^. A uniform rectilinear grid of 0.6 mm was used. The current between the electrodes was calculated integrating the current flux density on a closed surface surrounding one electrode and field magnitude were normalized to 2mA input current.

---

**Fig. S1.**
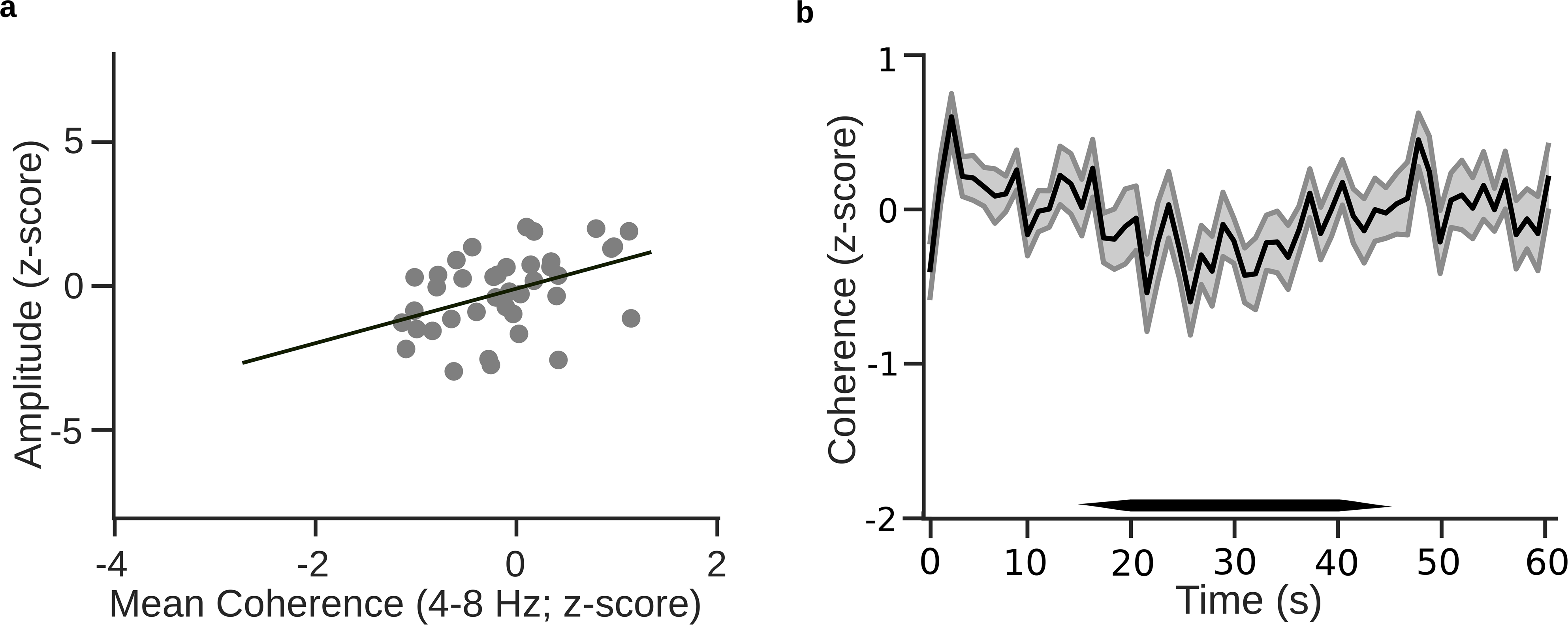
(related to Figure 5h) **a**, Correlation between change in tremor’s amplitude and change in tremor’s temporal coherence at the tremor frequency band (i.e., 4-8 Hz) of trials with stimulation without phaselocking. Each datapoint corresponds to the z-score of tremor’s amplitude versus the z-score tremor’s temporal coherence of a single trial during the stimulation period relative to the baseline period. c=-0.008, m=0.95, R^2^ =0.18 (c, line y-intercept; m, line slope). It shows that in the case of non-phase-locked stimulation, the change in the tremor amplitude was only weakly correlated (lower R^2^) with the change in the tremor temporal coherence and exhibited a weaker dependency (smaller line slope, i.e., coefficient of gradient) compared to phase-locked stimulation (Fig. 5hii). **b**, Change in tremor’s temporal coherence at the tremor frequency band over time during stimulation without phase-locking. Shown values are mean ± st.d. z-score at each epoch during stimulation period relative to the mean and st.d. across epochs during baseline period for the non-phase locking trials.

**Fig. S2.**
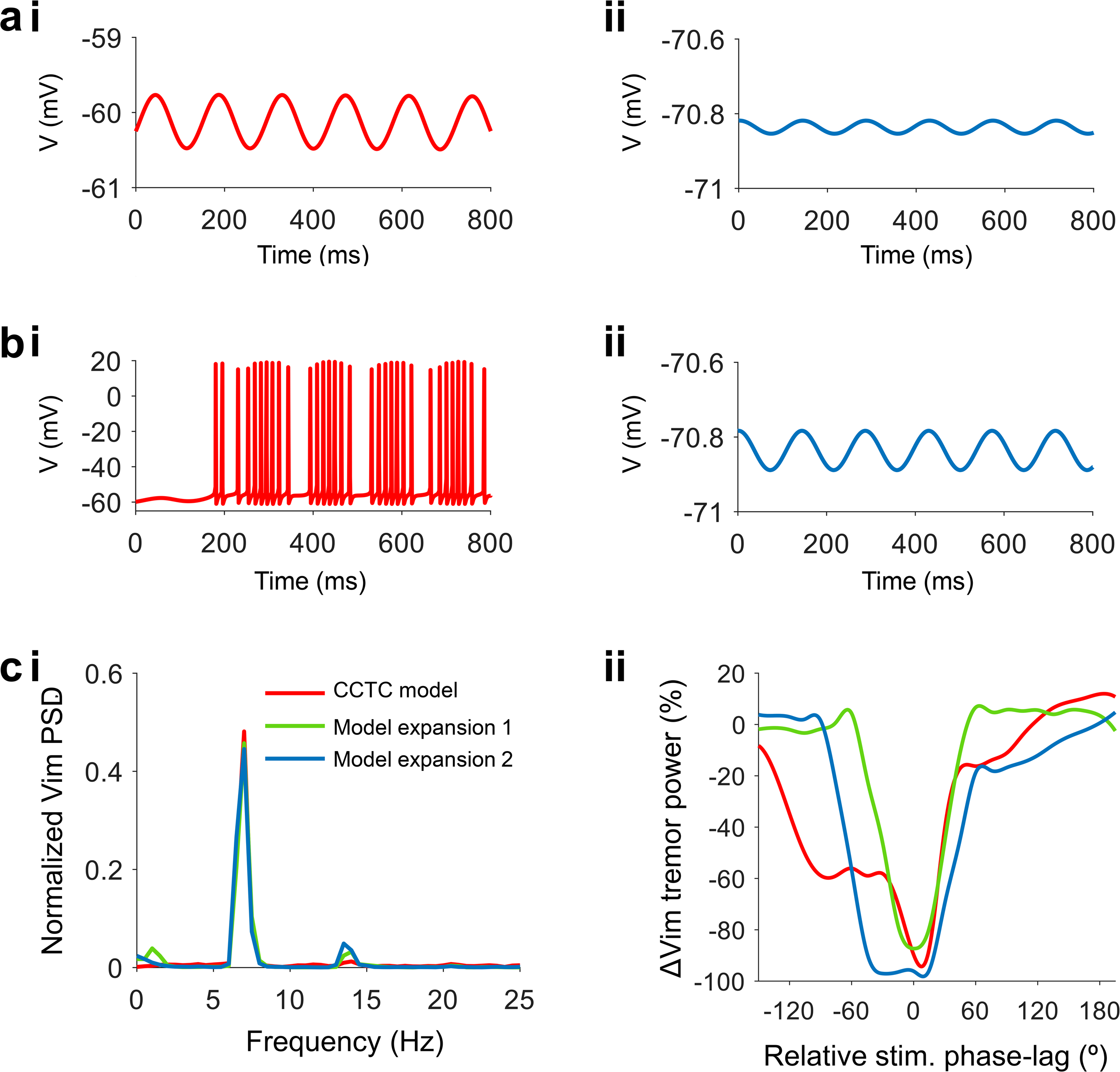
(related to Figure 6) **a-b**, Direct effect of the stimulation on different cell types in the cerebellar cortex. To identify the cell type in the cerebellar cortex that dominates the direct response to the applied electric fields, we simulated the response of the most abundant cell types in this region, i.e., the Purkinje cell (PC) and granule cell (GrC) to extracellular electric fields. To best capture the spatiotemporal dynamics, we used multi-compartmental models with 3D geometrical reconstruction of the PC^30^ and GrC^31^. We exposed the cells to homogenous extracellular electric fields that were aligned with the dendrite-somatic axes of the cells and quantified the induced depolarization. As in the original study with the PC model^30^, we removed the sodium and calcium channels from the axonal initial segment (AIS) of this cell to reduce its spontaneous pacemaker activity. **a**, Representative transmembrane potential traces of (i) PC and (ii) GrC during stimulation with an external sinusoidal electric field oscillating at the tremor frequency (i.e., 7Hz) and having an amplitude of 2 V/m expected in our experiment (Fig. 2b). It shows that the electrical stimulation induces a small periodic sub-threshold depolarization with magnitude that is ~20 folds larger in PCs compared to GrCs. **b**, Same as a but for electric field having an amplitude of 6 V/m. The results shown in (a-b) indicate that the direct response of the cerebellar cortex to the stimulating electric fields is dominated by the PCs. **c**, Representative spiking activity during stimulation without phase-locking at 16% normalized amplitude as in Fig 6g-j, showing (i) spike raster plot of PCs, (ii) spike raster of ION neurons, (iii) spike raster of DCNs, (iv) spiking spectrogram of TC neurons in Vim. **d**, Effect of the model size on the simulation outcome. To explore the effect of the model size on the simulation outcome, we repeated the simulation (‘CCTC model’, red line) with a higher number of cells, i.e., 5-fold increase, in the olivocerebellar circuit (‘Model expansion 1’, green line; synaptic connections between the DCNs and TC neurons of the Vim were randomized) and across the whole model (‘Model expansion 2’, blue line; synaptic connections between the DCNs and TC neurons of the Vim as well as between DCNs and PCs were randomized). Showing (i) normalized Vim PSD during stimulation with a relative phase of 0° and normalized amplitude of 16% and (ii) percentage change in tremor PSD in Vim versus stimulation phase for the same stimulation. It shows that in comparison to the original model, an increase in the number of cells in the model does not abolish the ET oscillation in the CCTC network (quantified via the PSD at the tremor frequencies in the Vim) and its response to phase-locked stimulation. However, it may modify range of efficacious phase-lags.

## Supplementary information

**Table S1.** (related to Fig. 2–5) Demographic and clinical information of patients. yrs, years; CRST, clinical rating scale for tremor.

| Patient No. | Sex | Handed-ness | Stim Hemi sphere | Age (yrs) | Age at Onset (yrs) | Disease Duration (yrs) | Tremor Severity (CRST) | Stimulation Intensity (mA) | Tremor Frequency<sup>1</sup> (Hz) | Tremor Amplitude Baseline<sup>1</sup> (a.u.) |
| --- | --- | --- | --- | --- | --- | --- | --- | --- | --- | --- |
| 1 | m | r | l | 65 | 5 | 60 | 48 | 2 | 5.2 (0.08, 0.9) | 704 (0.2, 09) |
| 2 | f | l | l | 48 | 45 | 3 | 14 | 4 | 6.9 (0.07, 1) | 88 (0.2, 1) |
| 3 | m | r | r | 80 | 55 | 25 | 16 | 2 | 6.2 (0.1, 0.9) | 35 (0.2, 0.9) |
| 4 | f | r | l | 79 | 69 | 10 | 29 | 2 | 5.9 (0.08, 0.9) | 151 (0.1, 1) |
| 5 | m | r | r | 71 | 25 | 46 | 44 | 1 | 4.6 (0.2, 0.6) | 1613 (0.2, 02) |
| 6 | m | r | r | 38 | 18 | 20 | 30 | 3 | 6.2 (0.09, 1) | 276 (0.2, 0.2) |
| 7 | m | r | r | 79 | 55 | 24 | 93 | 4 | 3.8 (0.09, 0.7) | 1454 (0.2, 0.9) |
| 8 | f | r | r | 78 | 57 | 21 | 57 | 3 | 5.4 (0.1, 0.9) | 480 (0.2, 0.9) |
| 9 | m | l | r | 52 | 1 | 51 | 57 | 2 | 6.2 (0.07, 0.9) | 204 (0.1, 0.2) |
| 10 | m | r | l | 82 | 64 | 18 | 63 | 3 | 5.2 (0.03, 0.9) | 79 (0.1, 0.8) |
| 11 | m | r | l | 70 | 55 | 15 | 37 | 3 | 4.1 (0.1, 1) | 76 (0.2, 0.8) |

<sup>1</sup> Showing in brackets patients' coefficient of variances (COVs; left) of tremor frequency and tremor amplitude during baseline period across trials and the corresponding p-values of the Hartigan's dip test for multimodality (right).

**Table S2.** (related to Fig. 2d) Demographic and clinical information statistics of all patient cohort (‘all patients’), patients who responded to phase-locking stimulation (‘responders’), and patients who did respond to phase-locking stimulation (‘non-responders’), as well as significant difference (i.e., p-value) between responders and non-responders, characterized using Wilcoxon rank-sum test.

|  | all patients | responders | non-responders | p-value |
| --- | --- | --- | --- | --- |
| Age (yrs) | 67.5 ±15 | 61.6 ±16 | 77.8 ±4.7 | 0.07 |
| Disease Duration (yrs) | 26.6 ±17.9 | 27.8 ±20.3 | 24.5 ±15.4 | 0.78 |
| Tremor Severity (CRST) | 44.4 ±23 | 37 ±18 | 57.2 ±28 | 0.29 |
| Stimulation Intensity (mA) | 2.6 ±0.9 | 2.8 ±0.9 | 2.5 ±1.3 | 0.8 |
| Tremor Frequency (Hz) | 5.4 ±1.0 | 6 ±1.1 | 4.9 ±0.9 | 0.08 |
| Tremor Amplitude Baseline (a.u.) | 468 ±594 | 205 ±237 | 825 ±823 | 0.23 |

Demographic and clinical information during repetition experiment with a subset of patients
| Patient No. | Age (yrs) | Disease Duration (yrs) | Tremor Severity (CRST) | Stimulation Intensity (mA) | Tremor Frequency (Hz) | Tremor Amplitude Baseline (a.u.) |
| --- | --- | --- | --- | --- | --- | --- |
| 1 | 68 | 63 | 75 | 2 | 4,9 | 974,0 |
| 2 | 51 | 6 | 15 | 3 | 6,5 | 43,0 |
| 3 | 84 | 29 | 47 | 2 | 6,4 | 53,0 |
| 6 | 42 | 24 | 31 | 3 | 6,8 | 82,0 |
| 9 | 55 | 54 | 47 | 2 | 6,9 | 118,0 |
| 11 | 73 | 18 | 46 | 3 | 4,8 | 46,0 |

**Table S3.** (related to Fig. 2e) Similarity of the mean phase lag between the stimulation conditions, assessed using Fisher test during the whole stimulation period (‘whole stim’), the 1st half of the stimulation period (‘1st stim half’), and the second half of the stimulation period (‘2nd stim half’). no-pl, no phase-locking.

| *Set phase lags* | *Whole stim.* | *1st stim half* | *2nd stim half* |
| --- | --- | --- | --- |
| 0° vs 60° | 2.73E-06 | 2.73E-06 | 2.73E-06 |
| 0° vs 120° | 2.73E-06 | 2.73E-06 | 2.73E-06 |
| 0° vs 180° | 0.20* | 0.20* | 0.20* |
| 0° vs 240° | 2.73E-06 | 2.73E-06 | 2.73E-06 |
| 0° vs 300° | 2.73E-06 | 2.73E-06 | 2.73E-06 |
| 60° vs 120° | 2.73E-06 | 2.73E-06 | 2.73E-06 |
| 60° vs 180° | 2.73E-06 | 2.73E-06 | 2.73E-06 |
| 60° vs 240° | 0.20 | 0.09 | 0.67 |
| 60° vs 300° | 2.73E-06 | 2.73E-06 | 2.73E-06 |
| 120° vs 180° | 2.73E-06 | 2.73E-06 | 2.73E-06 |
| 120° vs 240° | 2.73E-06 | 2.73E-06 | 2.73E-06 |
| 120° vs 300° | 0.39 | 0.67 | 0.01 |
| 180° vs 240° | 2.73E-06 | 2.73E-06 | 2.73E-06 |
| 180° vs 300° | 2.73E-06 | 2.73E-06 | 2.73E-06 |
| 240° vs 300° | 2.73E-06 | 2.73E-06 | 2.73E-06 |
| no-pl vs 0° | 0.09 | 0.67 | 0.20 |
| no-pl vs 60° | 0.67 | 0.20 | 0.67 |
| no-pl vs 120° | 0.20 | 0.67 | 1.00 |
| no-pl vs 180° | 0.03 | 0.67 | 0.20 |
| no-pl vs 240° | 0.67 | 0.09 | 0.67 |
| no-pl vs 300° | 0.20 | 0.67 | 0.67 |

\* We found that the Fisher test between 0° phase lag and 180° phase lag using MATLAB Circstats<sup>13</sup> function 'circ\_cmtest' is inaccurate due to the phase wrapping of the function.

**Table S4.** (related to Fig. 3i,k) Similarity of the circular spread of the phase lags, quantified via the mean resultant vector length *R*, between the stimulation conditions, assessed using ANOVA with post-hoc analysis using Wilcoxon signed-rank test during the whole stimulation period (‘whole stim’), the 1st half of the stimulation period (‘1st stim half’), the second half (‘2nd stim half’) of the stimulation period, and between the 1st and 2nd halves of the stimulation period (‘1st vs 2nd stim halves’). no-pl, no phase-locking.

| *Set phase lags* | *Whole stim*<br>*(p-value)* | *1st stim half*<br>*(p-value)* | *2nd stim half*<br>*(p-value)* |
| --- | --- | --- | --- |
| 0° vs 60° | 0.70 | 1.00 | 0.15 |
| 0° vs 120° | 0.64 | 0.58 | 0.52 |
| 0° vs 180° | 0.58 | 0.52 | 0.41 |
| 0° vs 240° | 0.24 | 0.37 | 0.15 |
| 0° vs 300° | 0.83 | 0.64 | 0.41 |
| 60° vs 120° | 0.24 | 0.37 | 0.58 |
| 60° vs 180° | 0.64 | 0.70 | 0.58 |
| 60° vs 240° | 0.70 | 0.37 | 0.90 |
| 60° vs 300° | 0.52 | 0.52 | 0.21 |
| 120° vs 180° | 0.12 | 0.07 | 1.00 |
| 120° vs 240° | 0.15 | 0.05 | 1.00 |
| 120° vs 300° | 0.90 | 0.17 | 0.52 |
| 180° vs 240° | 0.32 | 1.00 | 0.70 |
| 180° vs 300° | 0.52 | 0.90 | 0.46 |
| 240° vs 300° | 0.46 | 0.90 | 0.24 |
| no-pl vs 0° | 9.77E-04 | 9.77E-04 | 9.77E-04 |
| no-pl vs 60° | 9.77E-04 | 9.77E-04 | 9.77E-04 |
| no-pl vs 120° | 9.77E-04 | 9.77E-04 | 9.77E-04 |
| no-pl vs 180° | 9.77E-04 | 9.77E-04 | 9.77E-04 |
| no-pl vs 240° | 9.77E-04 | 9.77E-04 | 9.77E-04 |
| no-pl vs 300° | 9.77E-04 | 9.77E-04 | 9.77E-04 |
| sham vs 0° | 9.77E-04 | 9.77E-04 | 9.77E-04 |
| sham vs 60° | 9.77E-04 | 9.77E-04 | 9.77E-04 |
| sham vs 120° | 9.77E-04 | 9.77E-04 | 9.77E-04 |
| sham vs 180° | 9.77E-04 | 9.77E-04 | 9.77E-04 |
| sham vs 240° | 9.77E-04 | 9.77E-04 | 9.77E-04 |
| sham vs 300° | 9.77E-04 | 9.77E-04 | 9.77E-04 |
| sham vs no-pl | 0.37 | 0.27832 | 0.90 |

1st vs 2nd stim halves (p-value)
| Set<br>phase<br>lag 1st<br>stim<br>half/2nd<br>stim half | 0° | 60° | 120° | 180° | 240° | 300° | no-pl | sham |
| --- | --- | --- | --- | --- | --- | --- | --- | --- |
| 0° | 0.58 | 0.15 | 0.83 | 0.76 | 0.15 | 0.64 | 9.77E-04 | 9.77E-04 |
| 60° | 0.90 | 0.52 | 0.37 | 0.97 | 0.64 | 0.83 | 9.77E-04 | 9.77E-04 |
| 120° | 0.90 | 0.02 | 0.15 | 0.15 | 0.04 | 0.46 | 9.77E-04 | 9.77E-04 |
| 180° | 0.58 | 0.70 | 0.76 | 0.76 | 0.58 | 0.52 | 9.77E-04 | 9.77E-04 |
| 240° | 0.21 | 0.97 | 0.32 | 0.41 | 0.97 | 0.28 | 9.77E-04 | 9.77E-04 |
| 300° | 0.12 | 0.41 | 0.58 | 0.76 | 0.90 | 0.21 | 9.77E-04 | 9.77E-04 |
| no-pl | 9.77E-04 | 9.77E-04 | 9.77E-04 | 9.77E-04 | 9.77E-04 | 9.77E-04 | 0.90 | 0.83 |
| sham | 9.77E-04 | 9.77E-04 | 9.77E-04 | 9.77E-04 | 9.77E-04 | 9.77E-04 | 0.24 | 0.21 |

Change in tremor amplitude induced by phase-locked stimulation during the 1st half of the stimulation period (‘1st stim half’), the second half (‘2nd stim half’) of the stimulation period, and after the end of the stimulation (‘post stim’).

| Set phase lag | 1st stim half | 2nd stim half | post stim |
| --- | --- | --- | --- |
| 0° | -0.61 ± 0.38, p=0.14 | -0.86 ± 0.39, p=0.05 | -0.67 ± 0.37, p=0.10 |
| 60° | 0.09 ± 0.26, p=0.75 | -0.14 ± 0.26, p=0.60 | -0.06 ± 0.25, p=0.80 |
| 120° | -0.27 ± 0.39, p=0.51 | -0.54 ± 0.50, p=0.30 | -0.71 ± 0.35, p=0.07 |
| 180° | -0.26 ± 0.20, p=0.23 | -0.24 ± 0.37, p=0.53 | -0.68 ± 0.30, p=0.05 |
| 240° | -0.04 ± 0.24, p=0.87 | -0.08 ± 0.30, p=0.79 | -0.49 ± 0.42, p=0.27 |
| 300° | -0.39 ± 0.45, p=0.41 | -0.38 ± 0.39, p=0.35 | -0.36 ± 0.33, p=0.29 |

| Relative phase lag | 1st stim half | 2nd stim half | Post stim |
| --- | --- | --- | --- |
| -120° | -0.15 ± 0.08, p=0.56 | 0.18 ± 0.07, p=0.43 | -0.26 ± 0.08, p=0.37 |
| -60° | 0.18 ± 0.08, p=0.52 | -0.22 ± 0.07, p=0.33 | -0.27 ± 0.08, p=0.32 |
| 0° | -1.28 ± 0.13, p=0.02 | -1.43 ± 0.12, p=0.01 | -1.74 ± 0.10, p<0.0005 |
| 60° | -0.03 ± 0.07, p=0.89 | -0.11 ± 0.09, p=0.72 | -0.10 ± 0.07, p=0.67 |
| 120° | -0.39 ± 0.11, p=0.31 | -0.57 ± 0.17, p=0.35 | -0.23 ± 0.12, p=0.59 |
| 180° | 0.19 ± 0.07, p=0.40 | -0.10 ± 0.06, p=0.64 | -0.37 ± 0.08, p=0.20 |

Patients showing a statistically significant change in tremor amplitude due to phase-locked stimulation in the 2nd stim half. Showing patient number (‘ID’) and p-value of statistical testing comparing the phase-locked stimulation condition with sham stimulation (‘p-value Shm’). For subjects who also showed a significant reduction in tremor amplitude during nonphase-locked stimulation, the p-value of statistical testing comparing the phase-locked stimulation condition with the non-phase-locked stimulation condition (‘p-value non-pl’) is added.

| Set phase lag | Tremor decrease | Tremor increase |
| --- | --- | --- |
| 0° | 1 (3·10<sup>-4</sup>), 2 (3·10<sup>-3</sup>, 0.2), 3 (2·10<sup>-7</sup>), 4 (6·10<sup>-3</sup>), 6 (1·10<sup>-5</sup>) | 5 (7·10<sup>-3</sup>, 0.5) |
| 60° | 4 (3·10<sup>-4</sup>), 6 (1·10<sup>-3</sup>), 8 (5·10<sup>-4</sup>, 1·10<sup>-3</sup>), 9 (6·10<sup>-3</sup>), 10 (2·10<sup>-4</sup>, 0.9) | 5 (8·10<sup>-4</sup>, 0.3) |
| 120° | 2 (4·10<sup>-4</sup>, 0.06), 6 (1·10<sup>-3</sup>), 7 (4·10<sup>-5</sup>) | 4 (7·10<sup>-5</sup>), 5 (2·10<sup>-3</sup>, 1·10<sup>-3</sup>) |
| 180° | 2 (3·10<sup>-8</sup>, 1·10<sup>-5</sup>), 8 (1·10<sup>-4</sup>, 2·10<sup>-4</sup>) | 4 (4·10<sup>-3</sup>), 5 (1·10<sup>-5</sup>, 0.04) |
| 240° | 2 (4·10<sup>-4</sup>, 8·10<sup>-3</sup>), 8 (3·10<sup>-3</sup>, 0.08), 10 (2·10<sup>-5</sup>, 0.5) | 5 (1·10<sup>-7</sup>, 4·10<sup>-3</sup>), 7 (3·10<sup>-4</sup>, 0.2) |
| 300° | 2 (3·10<sup>-7</sup>, 3·10<sup>-7</sup>) | 10 (2·10<sup>-3</sup>) |

Number of patients showing a statistically significant change in tremor amplitude due to phase-locked stimulation during 2nd stim half, computed using Fisher exact test against the number of patients who did not show any change in tremor amplitude.

| Set phase lag | Tremor decrease<br>(n decrease, n no-change, p-value) | Tremor increase<br>(n increase, n no-change, p-value) |
| --- | --- | --- |
| 0° | 5, 5, p=1 | 1, 5, p=0.15 |
| 60° | 5, 5, p=1 | 1, 5, p=0.15 |
| 120° | 3, 6, p=0.39 | 2, 6, p=0.18 |
| 180° | 2, 7, p=0.08 | 2, 7, p=0.08 |
| 240° | 3, 6, p=0.39 | 2, 6, p=0.18 |
| 300° | 1, 9, p<0.005 | 1, 9, p<0.005 |

| *Set phase lag* | *Tremor decrease*<br>*(n decrease, n no-*<br>*change, p-value)* | *Tremor increase*<br>*(n increase, n no-*<br>*change, p-value)* |
| --- | --- | --- |
| -120° | 1, 9, p<0.005 | 1, 9, p<0.005 |
| -60° | 2, 7, p=0.08 | 2, 7, p=0.08 |
| 0° | 9, 1, p<0.005 | 1, 1, p=1 |
| 60° | 3, 6, p=0.39 | 2, 6, p=0.18 |
| 120° | 2, 8, p=0.03 | 1, 8, p=0.01 |
| 180° | 2, 7, p=0.08 | 2, 7, p=0.08 |

**Table S5.** (related to Fig. 3m-p) Similarity of the phase resultant between the original experiment and the repeated experiment (n=6, including patients 1,2,3,6, 9, and 11), assessed using a paired sign-rank test.

| *Set phase lag* | *original experiment* | *repeated experiment* | *difference* |
| --- | --- | --- | --- |
| 0° | 357° ± 13° | 2° ± 6° | p=0.76 |
| 60° | 60° ± 11° | 60° ± 7° | p=0.77 |
| 120° | 121° ± 9° | 110° ± 8° | p=0.19 |
| 180° | 182° ± 16° | 178° ± 6° | p=0.75 |
| 240° | 242° ± 3° | 237° ± 8° | p=0.37 |
| 300° | 299° ± 12° | 302° ± 9° | p=0.53 |

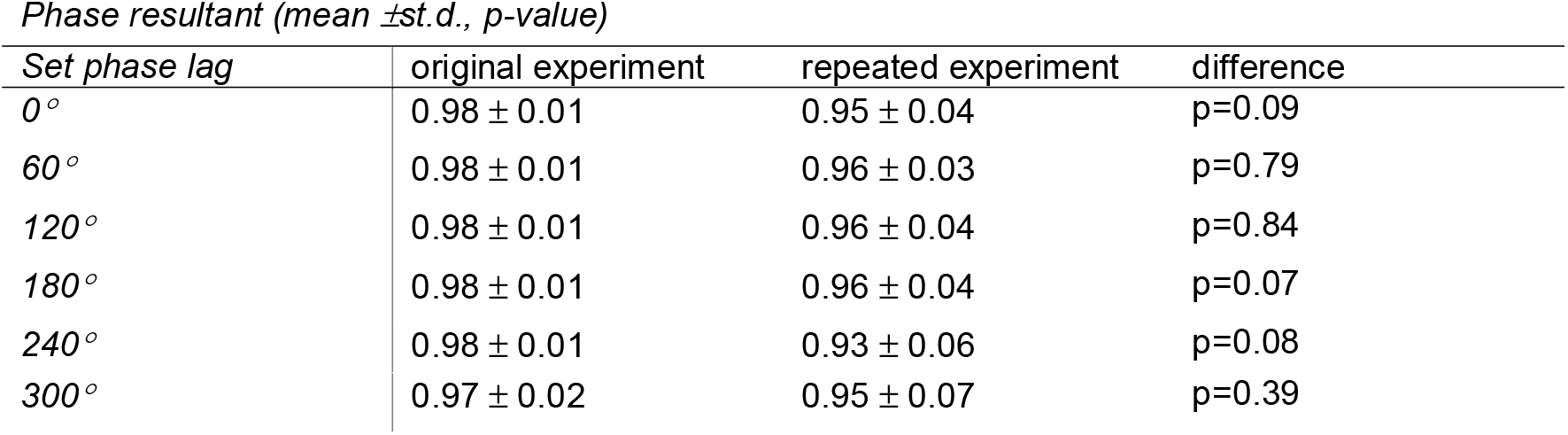
Similarity of the mean phase-lag between the original experiment and the repeated experiment (n=6, including patients 1,2,3,6, 9, and 11), assessed using Circular Krusical Wallis test.

| *Set phase lag* | *original experiment* | *repeated experiment* | *difference* |
| --- | --- | --- | --- |
| 0° | 0.98 ± 0.01 | 0.95 ± 0.04 | p=0.09 |
| 60° | 0.98 ± 0.01 | 0.96 ± 0.03 | p=0.79 |
| 120° | 0.98 ± 0.01 | 0.96 ± 0.04 | p=0.84 |
| 180° | 0.98 ± 0.01 | 0.96 ± 0.04 | p=0.07 |
| 240° | 0.98 ± 0.01 | 0.93 ± 0.06 | p=0.08 |
| 300° | 0.97 ± 0.02 | 0.95 ± 0.07 | p=0.39 |

**Table S6.** (related to Fig. 3o) Similarity of the change in tremor amplitude induced by phase-locked stimulation between the original experiment and the repeated experiment (n=6, including patients 1,2,3,6, 9, and 11), assessed using paired t-test. Showing, results during the 1st half of the stimulation period (‘1st stim half’), the second half (‘2nd stim half’) of the stimulation period, and after the end of the stimulation (‘post stim’).

| period | original experiment | repeated experiment | difference |
| --- | --- | --- | --- |
| 1st stim half | -1.86 ± 0.39, p=0.0001 | -1.82 ± 0.42, p=0.01 | p=0.93 |
| 2nd stim half | -2.10 ± 0.46, p=0.001 | -2.28 ± 0.53, p=0.001 | p=0.83 |
| Post stim | -1.81 ± 0.27, p=0.0001 | -1.62 ± 0.27, p<0.0001 | p=0.58 |

Patients showing a statistically significant change in tremor amplitude due to phase-locked stimulation between the original experiment and the repeated experiment (n=6, including patients 1,2,3,6, 9, and 11). Showing, results during the 1st half of the stimulation period (‘1st stim half’), the second half (‘2nd stim half’) of the stimulation period, and after the end of the stimulation (‘post stim’).

| Set phase lag | Tremor decrease<br>(n original experiment,<br>n repeated<br>experiment) | Tremor increase<br>(n original experiment,<br>n repeated<br>experiment) |
| --- | --- | --- |
| 1st stim half | 4, 4 | 1, 2 |
| 2nd stim half | 5, 5 | 0, 3 |
| Post stim | 6, 5 | 0, 2 |

**Table S7.** (related to Fig. 3p) Similarity of the change in tremor amplitude during stimulation between the original experiment and the repeated experiment (n=6, including patients 1,2,3,6, 9, and 11), assessed using paired t-test.

| Set phase lag | original experiment | repeated experiment | difference (p-value) |
| --- | --- | --- | --- |
| 0° | -1.46 ± 0.59 | -0.33 ± 0.56 | p<0.00001 |
| 60° | -0.20 ± 0.23 | -0.28 ± 0.46 | p=0.33 |
| 120° | -0.92 ± 0.80 | -0.94 ± 0.44 | p=0.10 |
| 180° | -0.32 ± 0.59 | -0.98 ± 0.70 | p=0.37 |
| 240° | -0.08 ± 0.31 | -0.88 ± 0.44 | p=0.10 |
| 300° | -0.91 ± 0.61 | -0.92 ± 1.19 | p=0.03 |

**Table S8.**
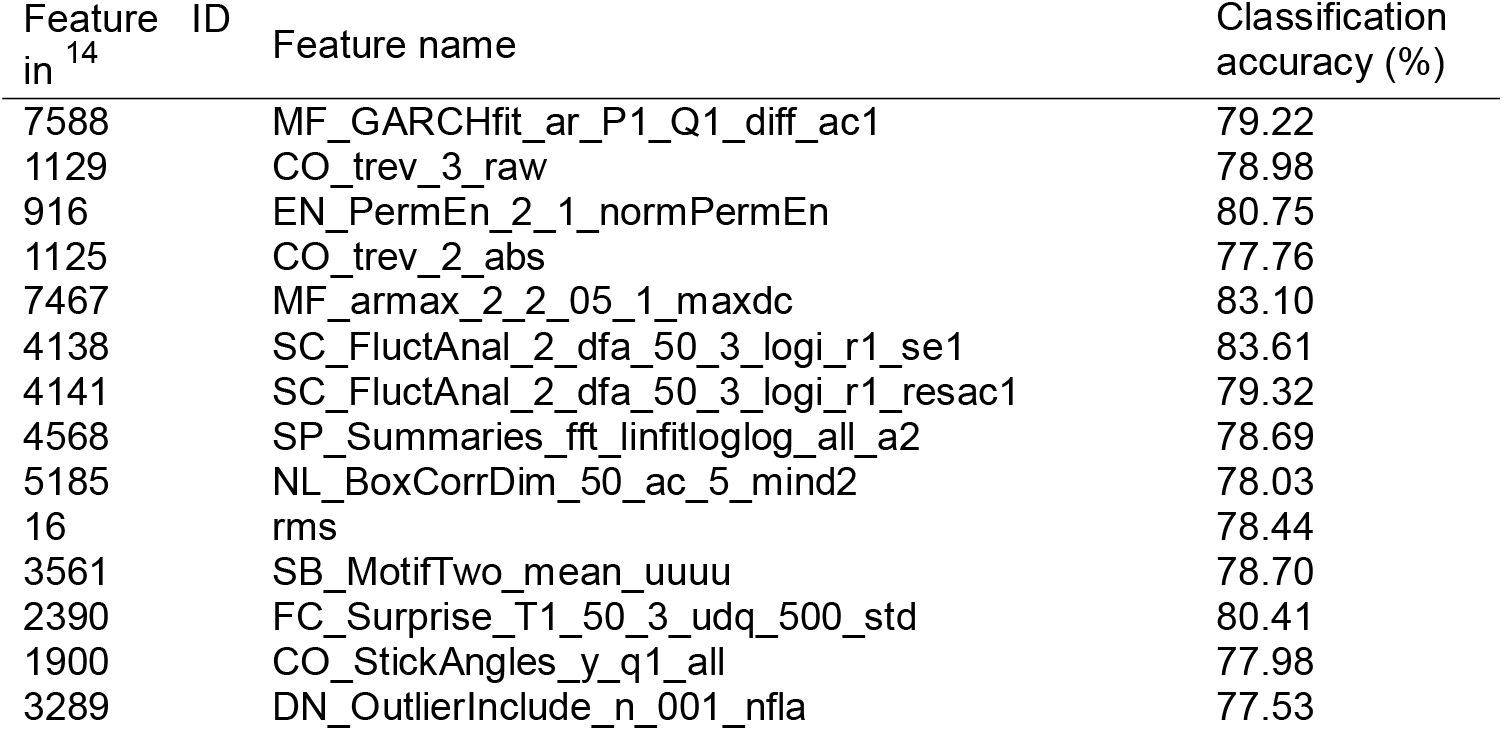
(related to Fig. 4c-d) Most informative features, i.e., the features shown in **Fig. 4c** at the centres of the clusters of correlated features, found to predict the patients’ response to stimulation. See Fulcher et al.^14^ for description of the features.

Description of features shown in Fig. 4d

- Normalised non-linear autocorrelation trev function – The function calculates the mean of the difference between the shifted time-series. A perfectly sinusoidal timeseries should have a zero difference in the means when shifted by tau. In our data, the non-responders had a more negative trev value which suggest that their movement was less symmetric.
- Root-mean square – The function computed the root mean square of the time series (i.e., amplitude). In our data, the responders had lower RMS.
- Motif correlation – The function searches for local motifs in a binary symbolisation of the time series with data points larger than mean set to 1 and those below the mean to 0. In our data, non-responders had larger amplitude above the mean, i.e., less symmetry relative to the time axis.
- Predictive memory of signal - The function estimates the surprise in the next data point given recent memory of the previous data points and then computes the st.d. of the surprise matrix. In our data, non-responders had a higher portion of extreme values compared to the mean information, thus more information in each data point.

See **Supplementary Software** for MATLAB code computing the abovementioned features.

**Table S9.** (related to Fig. 4g) Demographic and clinical information of second cohort of patients. yrs, years; CRST, clinical rating scale for tremor; a.u., arbitrary units.

| Patient No. | Sex | Handed-ness | Stimulated Hemisphere | Age (yrs) | Age at Onset (yrs) | Disease Duration (yrs) | Tremor Severity (CRST) | Stimulation Intensity (mA) | Tremor Frequency (Hz) | Tremor Amplitude Baseline (a.u.) |
| --- | --- | --- | --- | --- | --- | --- | --- | --- | --- | --- |
| 1 | m | r | r | 61 | 20 | 41 | 14 | 2 | 8.1 | 44.0 |
| 2 | f | r | r | 43 | 37 | 6 | 7 | 3 | 8.7 | 20.0 |
| 3 | m | r | r | 56 | 26 | 30 | 25 | 1 | 7.4 | 37.0 |
| 4 | m | l | l | 47 | 15 | 32 | 31 | 3 | 5.4 | 71.0 |
| 5 | f | r | r | 54 | 44 | 10 | 7 | 2 | 6.8 | 17.0 |
| 6 | f | r | l | 42 | 38 | 4 | 6 | 1 | 8.5 | 16.0 |
| 7 | f | r | l | 80 | 42 | 38 | 65 | 1 | 4.2 | 220.3 |

Demographic and clinical information statistics from the second cohort of patients, showing all patient cohort (‘all patients’), patients who responded to phase-locking stimulation (‘responders’), and patients who did respond to phase-locking stimulation (‘non-responders’), as well as significant difference (i.e., p-value) between responders and non-responders, characterized using Wilcoxon rank-sum test.

|  | all patients | responders | non-responders | p-value |
| --- | --- | --- | --- | --- |
| Age (yrs) | 55 ± 13 | 60 ± 19 | 51 ± 8 | 0.63 |
| Disease Duration (yrs) | 23 ± 16 | 25 ± 17 | 22 ± 18 | 1.0 |
| Tremor Severity (CRST) | 22 ± 21 | 32 ± 30 | 14 ± 12 | 0.46 |
| Stimulation Intensity (mA) | 0.2 ± 0.1 | 0.2 ± 0.1 | 0.2 ± 0.1 | 0.74 |
| Tremor Frequency (Hz) | 7.0 ± 1.7 | 6.8 ± 2.3 | 7.2 ± 1.4 | 1.0 |
| Tremor Amplitude Baseline (a.u.) | 60.8 ± 73.0 | 92.4 ± 111.1 | 37 ± 26.1 | 0.63 |

**Table S10.** (related to Fig. 4g) Phase-lag between stimulation and tremor movement in the second cohort of patients during the 1st half of the stimulation period (‘1st stim half’), the second half (‘2nd stim half’) of the stimulation period, and after the end of the stimulation (‘post stim’).

| Set phase lag | 1st stim half | 2nd stim half |
| --- | --- | --- |
| 0° | 7° ± 10°, p<0.00001 | 6° ± 12°, p<0.00001 |
| 60° | 59° ± 9°, p<0.00001 | 60° ± 8°, p<0.00001 |
| 120° | 113° ± 20°, p<0.00001 | 122° ± 14°, p<0.00001 |
| 180° | 180° ± 11°, p<0.00001 | 183° ± 12°, p<0.00001 |
| 240° | 247° ± 7°, p<0.00001 | 245° ± 9°, p<0.00001 |
| 300° | 308° ± 7°, p<0.00001 | 302° ± 7°, p<0.00001 |

Circular spread of the phase lags quantified via the mean resultant vector length *R* in the second cohort of patients.

Circular spread of the phase lags quantified via the mean resultant vector length $R$ in the second cohort of patients. *Phase resultant (mean ±st.d., p-value)*
| *Set phase lag* | 1st stim half | 2nd stim half |
| --- | --- | --- |
| 0° | 0.97 ± 0.03 | 0.96 ± 0.04 |
| 60° | 0.96 ± 0.03 | 0.97 ± 0.02 |
| 120° | 0.93 ± 0.14 | 0.94 ± 0.10 |
| 180° | 0.96 ± 0.05 | 0.97 ± 0.02 |
| 240° | 0.95 ± 0.07 | 0.97 ± 0.04 |
| 300° | 0.94 ± 0.08 | 0.95 ± 0.06 |

**Table S11.** (related to Fig. 4g) Patients in the second cohort, showing a statistically significant change (decrease or increase) in tremor amplitude during phase-locked stimulation and non phase-locked stimulation.

| *Set phase lag* | Phase-locked stimulation (n decrease, n increase) | Non phase-locked stimulation (n decrease, n increase) |
| --- | --- | --- |
| 1st stim half | 5, 2 | 3, 1 |
| 2nd stim half | 4, 4 | 1, 2 |
| Post stim | 5, 4 | 1, 2 |

Change in tremor amplitude induced by phase- vs phase of stimulation in the second cohort of patients.

Change in tremor amplitude induced by phase- vs phase of stimulation in the second cohort of patients. *Change in tremor amplitude vs stim phase lag (mean ±st.d. z-score, unpaired t-test p-value)*
| *Set phase lag* | 1st stim half | 2nd stim half | post stim |
| --- | --- | --- | --- |
| 0° | -0.12 ± 0.86, p=0.72 | -0.30 ± 2.31, p=0.74 | -0.40 ± 2.92, p=0.73 |
| 60° | -0.03 ± 0.86, p=0.94 | 0.24 ± 1.69, p=0.72 | -0.16 ± 1.79, p=0.82 |
| 120° | -0.19 ± 1.20, p=0.69 | -0.18 ± 1.20, p=0.70 | 0.06 ± 1.38, p=0.91 |
| 180° | -0.36 ± 0.93, p=0.35 | 0.02 ± 1.39, p=0.97 | -0.04 ± 1.52, p=0.95 |
| 240° | -0.37 ± 0.76, p=0.25 | -0.17 ± 1.07, p=0.70 | -0.36 ± 0.68, p=0.21 |
| 300° | -0.11 ± 0.73, p=0.70 | 0.15 ± 1.05, p=0.71 | 0.02 ± 1.24, p=0.96 |

**Table S12.**
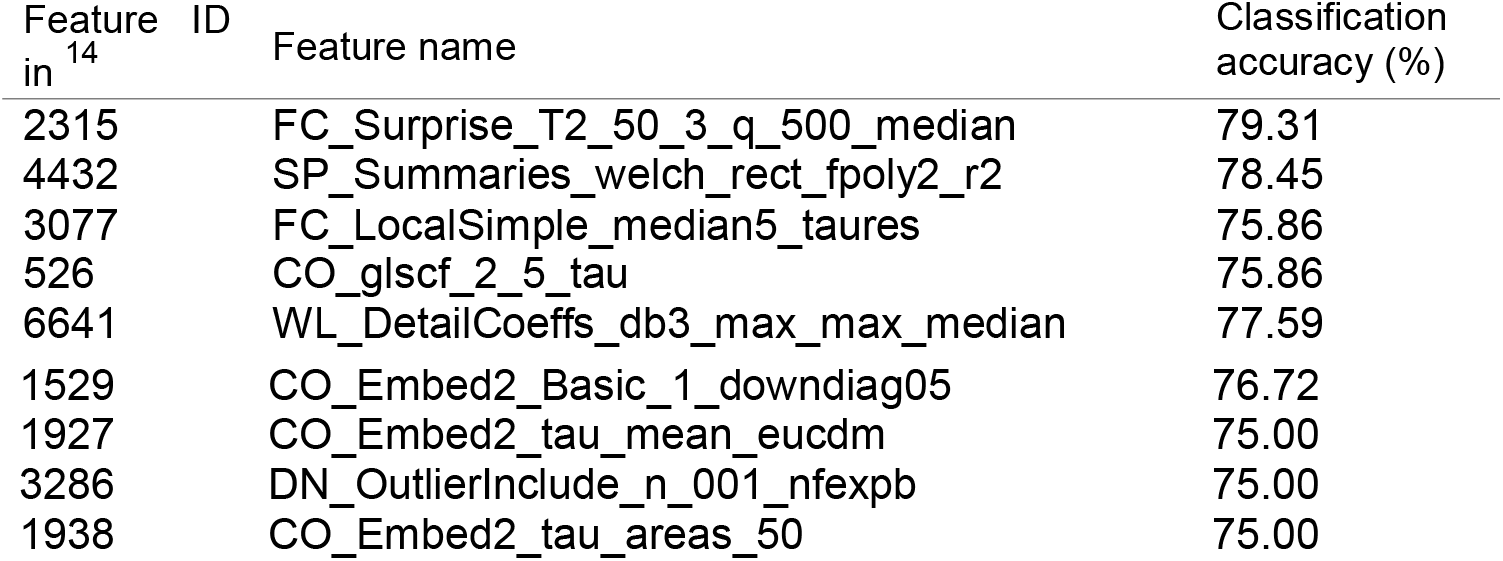
(related to Fig. 5e-f) Most informative features, i.e., the features shown in **Fig. 5e** at the centres of the clusters of correlated features, found to predict the patients’ response to stimulation. See Fulcher et al.^14^ for description of the features.

Description of features shown in **Fig. 5f**

- Information Gain – The function estimates the predictability of the next data point given the previous data points. A perfect sine wave has zero median information gain. In our data, stimulation that suppressed the tremor amplitude increased the information gain.
- Quadratic fit of power spectrum cumulative sum - The function computes the R^2^ of a quadratic fit to the cumulative sum of the power spectrum. In our data, stimulation that suppressed the tremor amplitude increase the accuracy of the fit, potentially by reducing the peak at the tremor frequency.

See **Supplementary Software** for MATLAB code computing the abovementioned features.

## Data Availability

The tremor data that support the findings of this study is available from the corresponding author upon request.

## Code availability

Matlab code of the ecHT is available as a **Supplementary Software**. The code for analysis of tremor measurements is available from the corresponding author upon request.

## Acknowledgments

**SRS** was supported by Swiss National Science Foundation, Swiss Neurological Society and European Academy of Neurology and EMDO Foundation. **XZ** was supported in part by the CT Institute for the Brain and Cognitive Sciences IBRAiN Fellowship. **SSan** was supported in part by the US NSF CAREER Award 1845348. **KPB**, was supported by Wellcome Trust MRC strategic neurodegenerative disease initiative award (WT089698), the Dystonia Coalition, Parkinson’s UK (G-1009). **MB** was supported by EPSRC award EP/N014529/1 funding the EPSRC Centre for Mathematics of Precision Healthcare. **ESB** acknowledges Lisa Yang, John Doerr, NIH R01MH117063, Edward and Kay Poitras. NG was funded by UK DRI Foundation Award, Wellcome Trust MIT fellowship (097443/Z/ 11/Z), Science & PINS Award for Neuromodulation, and NIHR IBRC Confident in Concept Award.

## Author information

### Contributions

**SRS**, designed and conducted clinical study, oversaw phase-locking, tremor amplitude and feature-based statistical learning analyses, wrote the paper. **DW**, developed ecHT, design and implemented ecHT-based phase-locking brain stimulator. **RP**, designed, developed and conducted feature-based statistical learning analysis. **JL**, developed and conducted phaselocking and tremor amplitude analyses. **ER and AL**, conducted the repeated clinical study. **EP**, helped developing ecHT theory. **ESB**, developed ecHT, oversaw experiments. **MB**, designed and oversaw feature-based statistical learning analysis. **XZ and SSan**, designed, developed and conducted neurophysiological simulation of ET. **KPB and JR**, designed and oversaw clinical study. **NG**, developed ecHT, designed and conducted clinical study, designed, developed and oversaw phase-locking and tremor amplitude analyses, oversaw feature-based statistical learning analysis and neurophysiological simulation of ET, and wrote paper.

